# BOLD signals of learning dynamics across and within trials beyond the classical fear and extinction network

**DOI:** 10.64898/2026.05.15.725431

**Authors:** Arslan Gabdulkhakov, Christian J. Merz, Fraenz Christoph, Erhan Genç

**Author notes:** **Corresponding Author and Lead Contact** Prof. Dr. Erhan Genç, Neuroimaging and Interindividual Differences Unit, Department of Psychology and Neurosciences, Leibniz Research Centre for Working Environment and Human Factors (IfADo), Ardeystraße, 67, 44139 Dortmund, Germany.

## Abstract

Human functional magnetic resonance imaging studies of fear conditioning often average neural responses across trials, potentially obscuring transient activations that vary across learning. In this study with 139 participants, we examined finer temporal dynamics of conditioned responding by analyzing three 2s segments within the conditioned stimulus (CS) presentation period across each quarter of fear acquisition and extinction training. This approach revealed distinct, time-specific engagement of regions within fear- and safety-related networks, both within and across trials. In particular, different activation patterns emerged across the three trial segments during CS presentation, indicating that neural responses were not limited to CS onset. We observed a more classical activation pattern at 0s relative to CS onset that diverged in later trial segments, most notably involving the amygdala, hippocampus, and prefrontal cortex (PFC) structures such as vmPFC present exclusively in 2s and 4s trial segments. We also found sustained activations consistent across all blocks of trials, such as right vlPFC activation 4s after CS onset across all fear acquisition quarters. These findings suggest that conditioned fear and safety processing unfold as dynamic spatiotemporal cascades and highlight the importance of modeling later responses following CS onset rather than focusing exclusively on onset-related activation.

## 1. Introduction

Anxiety- and trauma-related disorders present a significant global health burden, and their primary treatment relies on exposure-based therapies designed to promote fear extinction (Bie et al., 2024; Craske et al., 2014; Koenen et al., 2018). While often initially effective, a major clinical challenge is that extinguished fear frequently relapses when patients encounter previously feared stimuli in everyday environments outside the safe therapy setting (Bouton & Swartzentruber, 1991; Craske et al., 2014; Gunther et al., 1998; Meir Drexler et al., 2018; Vansteenwegen et al., 2007). Understanding the underlying mechanisms that drive this context-dependent return of fear is crucial for improving long-term therapeutic outcomes (Bouton, 2002, 2004; Pittig et al., 2018). In this study, we ask whether a time-resolved analysis of fear acquisition and extinction can reveal transient, trial-segment-specific neural activations that are not captured by conventional whole-phase analyses.

Associative learning is widely considered a fundamental adaptation mechanism that supports survival across species by allowing organisms to predict biologically significant events and adjust behavior accordingly, and in turn, increase the chances of survival (Pontes et al., 2020). Equally important is the ability to detect and learn changes in stimulus–outcome contingencies and to generalize learning appropriately beyond the original learning context (Pittig et al., 2018). However, abnormal learning and generalization of fear can contribute to the development of maladaptive behaviors that cause significant distress and dysfunction (Lissek et al., 2014; Pittig et al., 2018). Pavlovian fear conditioning provides a central experimental framework for probing how maladaptive fear responses are acquired and, crucially, how they can be reduced through processes such as extinction learning (Bouton, 2002; Pavlov, 1927; Wolpe, 1958). In a typical differential fear conditioning paradigm in humans, one cue is repeatedly paired with an aversive unconditioned stimulus (US) and becomes a conditioned stimulus (CS+) that elicits a conditioned fear response (CR), whereas a second cue (CS−) remains unpaired and functions as a safety signal. This fear acquisition training is used to model the development of aversive associations in a controlled setting. In a subsequent phase, fear extinction training is operationalized as repeated presentations of the CS+ without the US along with still safe CS– trials. Extinction learning is known to be strongly context-dependent, such that changes in external or internal contexts can bias retrieval toward either the original CS/US association or the newly learned inhibitory CS/noUS association (Bouton, 2002, 2004). In clinical practice, CS/US associations are often acquired in everyday environments but treated in a distinct therapy setting, so exposure (i.e., extinction) typically occurs in a context B that differs from the original acquisition context A, which is thought to promote return of fear when patients re-encounter the CS outside the therapy context (Craske et al., 2014; Dunsmoor et al., 2014; Gunther et al., 1998; Vansteenwegen et al., 2007).

Contemporary neuroimaging meta-analyses have delineated a distributed network consistently engaged during fear and extinction learning, encompassing the amygdala, hippocampus, anterior insular cortex (AIC), dorsal anterior cingulate cortex (dACC), and ventromedial prefrontal cortex (vmPFC; Fullana et al., 2016, 2018; Radua et al., 2025; Wen et al., 2024). These regions are known to constitute a so-called “fear and safety” network. However, the view that individual structures are involved more in fear and/or extinction learning is being challenged and re-evaluated in the contemporary fear conditioning literature. For example, Battaglia et al. (2020) showed that a lesion to the vmPFC, which was previously primarily attributed to fear extinction, reversal, or extinction recall, impairs successful fear learning. A meta-analysis by Fullana et al. (2016) identified large-scale brain activation across the so-called “fear and extinction” network and demonstrated consistent activation patterns across different conditioning paradigms. However, they did not report significant activations in the amygdala or vmPFC, raising questions about the functional involvement of these regions in human fear learning. One possible explanation is that their analysis averaged activation across the entire fear acquisition and extinction training phases, focusing on the differential CS+ versus CS– contrast on a “global” level. Such an approach may obscure transient or phase-specific effects occuring only during specific timepoints or even trial segments of training.

To address the limitations of trial-averaged analyses, an increasing number of electrophysiology and neuroimaging studies have partitioned the experimental data into shorter temporal bins, such as trial-by-trial or discrete blocks of trials, to capture the dynamic evolution of fear and extinction learning (for example, Harnett et al., 2016; Jentsch et al., 2020; Labrenz et al., 2022; Merz et al., 2018; Sperl et al., 2021; Wen et al., 2022, 2024). For instance, Wen et al. (2022) examined the temporally dynamic engagement of amygdaloid subnuclei, specifically, the basolateral amygdala (BLA) and centromedial amygdala (CMA), using a trial-by-trial blood-oxygen-level-dependent (BOLD) time-course analysis. They observed stronger CS+□>□CS–activation in both BLA and CMA during early acquisition, followed by a reversal of this activation (CS–□>□CS+) in the BLA during late acquisition. These findings underscore the temporal specificity and the transient nature of the amygdala activation, and show the importance of studying these phenomena at temporally resolved scales, as averaging across the entire acquisition or extinction training could result in weakening significant activations. In their subsequent work with a large dataset and multivoxel pattern analysis (MVPA), Wen et al. (2024) found that regions beyond the classical “fear and safety” network contributed to higher decoding accuracy, highlighting the importance of examining broader neural systems involved in fear learning. Second, by splitting the experiment into 4 blocks of trials (referred to as quarters Q1-Q4 below), they demonstrate that the dACC, dorsal anterior insula, posterior cingulate cortex, opercular part of the inferior frontal cortex, caudate nucleus, and thalamus consistently encode the CS+ in comparison to CS- across fear acquisition and extinction training as well as recall. Conversely, they found that activations in the posterior and anterior parts of the hippocampus, orbital frontal cortex, and the posterior insular cortex consistently coded the CS– in comparison to CS+ across phases. Beyond these, they identified a broad network, including limbic structures (BLA, CMA), prefrontal regions (vmPFC, subgenual ACC), and various sensory-motor and cerebellar areas, that exhibited dynamic signal shifts between CS+ and CS– across phases. This temporal variability likely explains why these regions were frequently absent in earlier trial-averaged reports. As suggested by Andres et al. (2024), this variability likely reflects the transient nature of extinction-related neural responses, which may occur predominantly within the first few trials or even evolve dynamically within a single trial (Miskovic & Keil, 2012; Sperl et al., 2021).

Complementing neuroimaging work, a recent EEG study by Starita et al. (2023) revealed distinct oscillatory dynamics associated with fear and reversal learning with three 2-s-long trial segments at 0 to 2 s, 2 to 4 s, and 4 to 6 s relative to the CS onset. They observed a strong differential CS+/CS– alpha effect source localized in the motor cortex, and the theta effect source localized in the midcingulate cortex, which was ramping up across the three trial segments towards the end of the trial. Building on these findings, our recent simultaneous EEG–fMRI study (Gabdulkhakov et al., 2026) replicated the ramping up theta effects reported by Starita et al. (2023) and extended them by identifying the cortical targets of frontal midline theta modulation. Specifically, we observed theta-phase–related modulation of cuneal, sensory, and motor cortices at 2 to 4 s and 4 to 5.5 s relative to CS onset (CS+□>□CS–) during fear acquisition training. During extinction training, we additionally found a theta-modulated BOLD activation in the left precentral gyrus within 0 to 2 s (CS–□>□CS+) and vmPFC in the 2 to 4 s window (CS+□>□CS–). Together, these results reveal selective and temporally specific activation patterns at different time points, even within one trial. However, despite looking at the segments within the trials, Gabdulkhakov et al. (2026) and Starita et al. (2023) focused on averaging the signal across all the trials during the respective learning phases, due to the limitation of the EEG requiring an average of multiple trials, thus missing the dynamics during the phases.

As described above, the majority of previous neuroimaging studies have typically analyzed entire fear acquisition or extinction phases, or, when temporally subdivided, have focused on early post-CS intervals in event-related designs or the full CS duration in block designs. Such approaches risk averaging out transient or dynamic activations, thereby obscuring the timing and sequence of neural processes. This could create the misleading impression that certain regions remain continuously active across long durations, whereas temporally resolved analyses suggest that fear-related activation fluctuates dynamically within trials (e.g., Sperl et al., 2021; Starita et al., 2023) and across trials (e.g., Wen et al., 2022, 2024).

In the present work, we address this limitation by splitting the trials of fear acquisition and extinction training into quarters and then splitting each quarter into three trial segments at 0 s, 2 s and 4 s relative to the CS onset. With a relatively large fMRI dataset (N = 139), our goal is to expand the known literature threefold. First, we aim to disentangle and attribute the known structures of the so-called “fear and extinction” network to specific trial segments and quarters of trials during fear acquisition and extinction training. Second, we expect that certain structures transiently involved in fear and extinction learning, which might have been averaged out in more conventional analyses, would show significant activation at narrower time scales. Lastly, we aim to observe structures at much later trial segments relative to the CS onset, not typical for conventional fMRI analyses with smaller samples, or big samples composed from datasets with different paradigms and thus different trial durations. Knowing which structures are involved specifically during fear extinction learning could inform future studies and treatment to facilitate extinction recall, and thus retain the safer CS/no US contingency.

## 2. Methods

### 2.1 Participants

The target sample size was informed by the range of sample sizes typically employed in the fear conditioning literature, while accounting for anticipated exclusions. A total of N = 139 participants (83 women, 56 men) aged between 18 and 26 (*M* = 21.99, *SD* = 2.08) years were recruited for this study. Following Lakens (2022), we performed a sensitivity analysis in MorePower 6.0 (Campbell & Thompson, 2012) to establish the minimum detectable effect size for our final sample sizes and repeated-measures design (2 CS types × 4 quarters of trials). For *N* = 137 (acquisition) and *N* = 138 (extinction), at 80% power and α = .05, this yielded sufficient sensitivity to detect partial η*²* ≥ .03.

Exclusion criteria at the recruitment stage comprised a history of mental health conditions, substance or psychoactive medication use, and standard exclusion criteria for fMRI examinations at a 3T scanner. All participants had normal or corrected-to-normal vision and were able to understand the provided written and oral instructions. All participants were naive to the purpose of the study and had no prior experience with the fear conditioning paradigm used for the experiment. The study was approved by the local ethics committee of the Faculty of Psychology at Ruhr University Bochum (application number 327). All participants provided written informed consent before participation and were treated in accordance with the Declaration of Helsinki.

### 2.2 Experiment

The study was conducted at the Bergmannsheil Hospital in Bochum, Germany, using a 3T Philips Achieva MRI system (Philips Healthcare, Best, The Netherlands) equipped with a 32-channel head coil. Briefly, participants completed both fear acquisition and extinction training within the same scanning session, separated by an 8-minute rest phase. The protocol and visual stimuli were based on the fear conditioning design developed by Milad et al. (2007). Visual cues were displayed on an MR-compatible LCD screen (BOLDscreen 24; Cambridge Research Systems, Cambridge, UK) positioned behind the scanner bore. Participants were instructed to attentively observe the visual stimuli throughout fear acquisition and extinction training and were informed that mild electrical stimulations could occur during the experiment.

Fear acquisition training comprised 32 total trials (16 CS+ and 16 CS−), with the CS+ paired with the unconditioned stimulus (US) in 10 of its 16 trials (reinforcement rate = 62.5%). After that, extinction training started, consisting of 8 unreinforced CS+ and 8 CS− presentations (see Figure 1). The number of trials was selected to align with widely used human fear conditioning paradigms, promoting methodological comparability with prior work (e.g., Milad et al., 2007; Phelps et al., 2004). Each trial began with a 1 s context cue (office lamp on a desk during fear acquisition training, lamp on a bookshelf during extinction training), followed by a 6-second CS, represented by the lamp illuminated in either blue or yellow.

**Figure 1.**
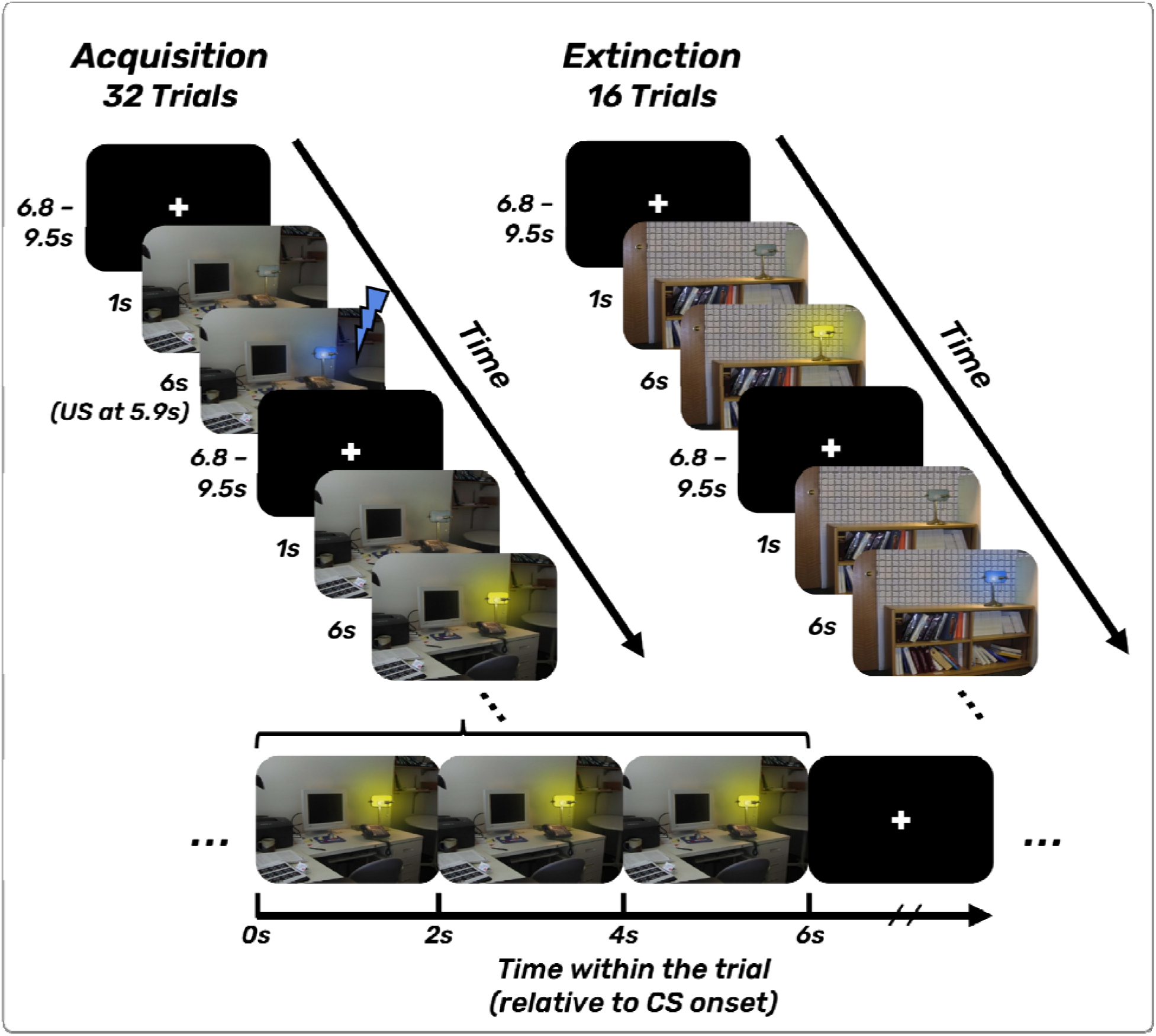
Experimental outline of fear acquisition (upper left panel) and extinction (upper right panel) training, adapted from Gabdulkhakov et al., (2026). The lower panel represents the critical timepoints within the trial modelled with the GLM at 0 s, 2 s, and 4 s relative to the CS onset. Both fear acquisition and extinction training started with a fixation cross, which was also presented during the ITI (jittered between 6.8 and 9.5 s). Following the ITI, a context image (an office table during fear acquisition training - Context A; a bookshelf during extinction training - Context B) was presented for 1 s. A lamp in the context image then lit up (blue or yellow) for 6 s, serving as either the CS+ or CS−. During fear acquisition training, the US (a 100 ms electrical stimulation to the right hand) was delivered at 5.9 s post-CS+ onset and co-terminated with the CS+. A 62.5% partial reinforcement schedule was used, with 10 of 16 CS+ trials paired with the US. CS = conditioned stimulus; US = unconditioned stimulus; ITI = intertrial interval; GLM = general linear model.

The US consisted of a train of 1 ms electrical pulses (50 Hz, 100 ms duration) delivered by a constant-voltage stimulator (STM200; BIOPAC Systems, Goleta, CA, USA) through two fingertip electrodes placed on the right index and middle fingers. The US onset occurred 5.9 s after CS presentation onset and co-terminated with the CS+. Before scanning, shock intensity was individually calibrated to be “very unpleasant but not painful,” beginning at 30 V and increasing in 5 V increments. Trial order was pseudo-randomized such that no more than two consecutive trials of the same type occurred.

Prior to scanning, participants were informed that both visual images and potential electrical stimulations would be presented. After completing the task, they answered a short questionnaire assessing their perceived number of stimulations and contingency awareness. Specifically, participants estimated (a) the total number of shocks received and (b) the percentage of times an electrical stimulation accompanied each CS (blue and yellow lamps; response scale: 0–100% in 10% steps).

### 2.3 Contingency ratings

Before scanning, all participants were informed that they would be presented with images and may or may not receive electrical stimulations during the experiment. After completion of fear acquisition and extinction training, participants filled out a contingency and stimulation rating questionnaire. First, they were asked: ‘How many electrical stimulations do you think you received in total during the experiment?’ and reported a number. Second, for each CS (blue lamp, yellow lamp), they answered the questions: ‘On what percentage of blue lamp presentations was an electrical stimulation delivered?’ and ‘On what percentage of yellow lamp presentations was an electrical stimulation delivered?’ (0–100% in steps of 10%). These scores, including descriptive statistics (mean and standard deviation) and the results of paired t-tests (Pingouin Package; Vallat, 2018) comparing CS+ and CS− contingency ratings, are reported in the Results section.

### 2.4 Skin conductance response recording and analysis

Skin conductance responses (SCRs) were acquired via two Ag/AgCl electrodes, each filled with an isotonic 0.05 M NaCl solution, positioned on the hypothenar eminence of the participants’ left hand. Data were recorded using Brain Vision Recorder software (version 1.23.0003; Brain Products, Munich, Germany) at a sampling rate of 5,000 Hz. Scanner-related artefacts were removed in Brain Vision Analyzer (version 2.2; Brain Products, Munich, Germany), after which the cleaned signals were exported to MATLAB (R2022b; MathWorks, Natick, MA, USA) for additional preprocessing performed in the EDA-Analysis App (Otto et al., 2023). The SCR amplitudes were defined as the maximal change in skin conductance identified through a foot-to-peak analysis within a 1 to 6.5□s window following CS onset. The extended window was chosen to capture anticipatory responses across the entire CS presentation period and to align with the later trial segments analyzed in fMRI. Although this deviates from the commonly used 1 to 4□s interval recommended by Boucsein et□al.□(2012), a recent systematic comparison by Kuhn et□al.□(2022) found no consistent differences in SCR results across preprocessing strategies using different time windows. They reported that longer windows covering the entire CS presentation perform similarly to narrower ones and that no universally optimal latency window exists. Our decision was therefore guided by the specific temporal structure of our task. Following standard recommendations (Boucsein et al., 2012), only responses exceeding 0.01 µS were retained as valid reactions; smaller fluctuations were coded as zero to represent non-responses but were still included in subsequent analyses to maintain the full range of individual variability. No further normalization procedures, such as range correction or z-transformation, were applied. This approach was selected to preserve meaningful interindividual differences in electrodermal reactivity that could otherwise be obscured by standardization.

For each participant, the CR was computed as the difference between the mean SCR amplitudes to CS+ and CS− trials. The processed SCR dataset was then exported for statistical analysis. Participants exhibiting excessive residual scanner artefacts or a poor signal-to-noise ratio were excluded to ensure data quality, yielding a final sample of N = 121.

To test the CR for significance and to ensure comparability with both the prior literature and our fMRI analysis (see section 2.5), we ran three analyses in order of increasing temporal resolution. First, to assess overall conditioned responding across the full phase, we used a one-way within-subjects ANOVA with CS type (CS+, CS−) as the single factor. Second, to examine coarse temporal dynamics, we used a two-way within-subjects ANOVA with CS type (CS+, CS−) and Half (Early, Late) as factors, where Early corresponded to trials 1–8 (acquisition) and 1–4 (extinction), and Late to trials 9–16 (acquisition) and 5–8 (extinction). Third, to capture fine-grained temporal evolution, we used a 2 (CS: CS+, CS−) × 4 (Quarter: Q1–Q4) within-subjects ANOVA, where quarters for fear acquisition training were defined as Q1 (trials 1–4), Q2 (trials 5–8), Q3 (trials 9–12), and Q4 (trials 13–16), and for extinction training as Q1 (trials 1–2), Q2 (trials 3–4), Q3 (trials 5–6), and Q4 (trials 7–8). All ANOVAs were implemented in the Pingouin Python package (Vallat, 2018). We assessed sphericity with Mauchly’s test and reported Greenhouse-Geisser corrected degrees of freedom and *p*-values where this assumption was violated. ANOVA results are reported with *F*-statistics, degrees of freedom, *p*-values, partial η*²* effect sizes, and their 90% confidence intervals (Lakens, 2013). For statistically significant main effects (*p* < .05), Bonferroni-corrected post hoc tests were conducted, with results reported as means, standard deviations, and 95% CIs of the mean difference.

For comparability with previous literature and with our fMRI analysis (see section 2.5 below), we also assessed CR at broader temporal scales. We examined early (Q1 + Q2) vs. late (Q3 + Q4) learning phases using a 2 (CS: CS+, CS−) × 2 (Halves: Early, Late) ANOVA. Finally, to test CR across the complete phase, SCRs were averaged across all quarters for each participant, and a one-way within-subjects ANOVA (CS Type: CS+ vs. CS−) was conducted. Note that for the Halves and Full Phase analyses, the within-subjects factors contained only two levels; thus, the sphericity assumption was satisfied by definition.

### 2.5 (f)MRI recording and analysis

High-resolution anatomical images were collected using a T1-weighted MP-RAGE sequence (8.2 ms TR, 3.7 ms TE, 8° flip angle, 220 slices, 240 mm × 240 mm matrix, 1 mm³ isotropic voxels) over approximately 6 minutes of scanning time. Functional MRI data during fear acquisition and extinction training were acquired with an echo planar imaging sequence (2,500 ms TR, 35 ms TE, 90° flip angle, 40 slices, 112 mm × 112 mm matrix, 2 mm × 2 mm × 3 mm resolution). Fear acquisition training lasted roughly 8 minutes, while extinction training took approximately 4 minutes.

Functional data underwent preprocessing in FSL FEAT (v6.0.1; Jenkinson et al., 2011), which included motion correction via MCFLIRT, slice timing correction, spatial smoothing with a 6 mm FWHM Gaussian kernel, and high-pass temporal filtering (50 s cutoff). The 6 mm smoothing kernel was selected to balance signal-to-noise enhancement with spatial precision while accommodating inter-individual anatomical differences in group-level analyses (Mikl et al., 2008). Registration proceeded in two stages: functional images were first linearly aligned to each participant’s anatomical scan, then transformed to MNI152 standard space using a 12-parameter affine registration. Two participants were excluded due to excessive motion artifacts or insufficient brain coverage.

For fear acquisition training, we modelled the onset times of CS+ and CS– trials from 1 through 4 (Q1), 5 through 8 (Q2), 9 through 12 (Q3), 13 through 16 (Q4), as well as the onset of context presentation, the US delivery time points after the reinforced CS+ trials, and analogous time points after the nonreinforced CS+ as well as CS– trials. Similarly, for extinction training, we modelled the onset times of CS+ and CS– trials 1 and 2 (Q1), 3 and 4 (Q2), 5 and 6 (Q3), 7 and 8 (Q4), as well as the onset of context presentation, and the time point of the offset of both CS. We then contrasted CS+ > CS– and CS– > CS+ within each corresponding quarter as the regressors in the General Linear Model (GLM; for example, (CS+ in Q1) > (CS– in Q1)). Additionally, we grouped the quarters together to model the halves as well as the complete duration of fear acquisition and extinction training. For example, for the analysis of halves, the model would be a composite of ((CS+ in Q1) + (CS+ in Q2)) > ((CS– in Q1) + (CS– in Q2)). Similarly, for the analysis of the full duration of the phases, we composed the model from all quarters for CS+ and contrasted it against all quarters in CS–.

To model the different trial segments, the CS+ and CS– regressor onset times were temporally shifted to model the periods at 0 s, 2 s, and 4 s relative to the actual CS onset. Analysis for each trial segment was run separately on the first and second levels, but with the identical preprocessing procedures and contrasts described above. We used separate analyses to study the effects at different temporal proximities to CS onset, as opposed to having a partial trial design (e.g., Ruge et al., 2009), on the premise that distinct processes dominate at each trial segment. This can range from early visual encoding of the CS, through retrieval and updating of the learned CS–US association, to preparatory responding, as suggested by temporal analyses of fear conditioning-related sensory and associative responses (Jentsch et al., 2020; Harnett et al., 2016). Additionally, running all regressors in a single GLM poses a risk of raising collinearity among the regressors. The second-level (group level) models were analyzed with FLAME 1, applying cluster thresholding with a Z-statistic threshold of 3.1 (Eklund et al., 2016) and *p* < .05. The significant clusters were mapped to brain areas using the Julich Histological Atlas and the Harvard-Oxford Cortical and Subcortical Atlases, as implemented in FSL. The significant statistical maps are visualized with nilearn Python package (version 0.10.2; nilearn contributors, 2023).

To enhance the comparability of our findings with established blueprints of fear and extinction learning, we assessed the spatial overlap between our statistical maps and those previously reported by Fullana et al. (2016, 2018) and Wen et al. (2024). First, using nilearn, all statistical maps were resampled to a 2 mm MNI152 template space using nearest-neighbor interpolation. Next, we created combined binary masks representing the union of the Fullana and Wen maps for the corresponding learning phases. We then compared our thresholded statistical maps (Z > 3.1) against these combined masks to identify regions of overlap, regions unique to our current analysis, and regions uniquely present in the combined prior literature. For fear acquisition training, we aligned our analysis with the trial blocks reported by Wen et al. (2024), comparing our Q1 data to a combined mask of Fullana and Wen’s block 1, our Q2 to their block 2, and so forth. During extinction training, because Wen et al. (2024) only reported results for the first block, we compared all four of our extinction quarters against a single combined mask derived from a union of Fullana’s overall extinction map and Wen’s first extinction block. Crucially, while the Fullana and Wen maps reflect standard, single-time-point trial analyses, our models incorporated temporal shifts (0 s, 2 s, and 4 s relative to CS onset). Therefore, for each learning phase and block, the same static combined mask from the prior literature was used as a consistent reference to evaluate overlap and unique activation across each of our three novel temporal segments. The resulting overlaps and unique clusters were visualized on a glass brain using nilearn and are reported in Supplementary Figures S5-S12.

## 3. Results

### 3.1 Contingency ratings

The analysis of self-report scores obtained after fear acquisition training revealed successful fear learning. Participants reported receiving an average of 10.84 (*SD* = 4.04) electrical stimulations, aligning with the applied 10 stimulations over 16 CS+ trials. Likewise, the average reported percentage of electrical stimulations received after the CS+ (*M* = 57.74%, *SD* = 17.84) was significantly higher than after the CS− (*M* = 10.10%, *SD* = 17.87, *t*(135) = 11.32, *p* < .001, Cohen’s *d* = 2.67). The mean reported values closely approximated the actual reinforcement rates of 62.5% for CS+ and 0% for CS−. For extinction training, all participants correctly reported that they received 0 electrical stimulations.

### 3.2 Skin conductance responses (SCRs)

#### 3.2.1 SCR Across All Trials: (Full phase)

To evaluate overall conditioned responding across the full phase, a one-way repeated-measures ANOVA was conducted to examine the effects of **CS type** (CS+ vs. CS−) on the average SCR across all trials (Q1 through Q4) during fear acquisition and extinction training.

During fear acquisition training (See Figure 2A, left panel), the analysis revealed a significant main effect of **CS type**, *F*(1, 114) = 10.47, *p* = .001, partial η*²* = .08, 90% CI [.02, .17]. As expected, overall SCRs were significantly higher for the CS+ (*M* = 0.41, *SD* = 0.69) compared to the CS− (*M* = 0.25, *SD* = 0.28). During extinction training (See Figure 2A, right panel), a similar analysis revealed a significant main effect of **CS type**, *F*(1, 101) = 4.63, *p* = .03, partial η*²* = .04, 90% CI [.01, .12]. This shows that when all trials are averaged together, SCRs are still significantly higher for the CS+ (*M* = 0.34, *SD* = 0.44) compared to the CS− (*M* = 0.27, *SD* = 0.36) during extinction training.

**Figure 2.**
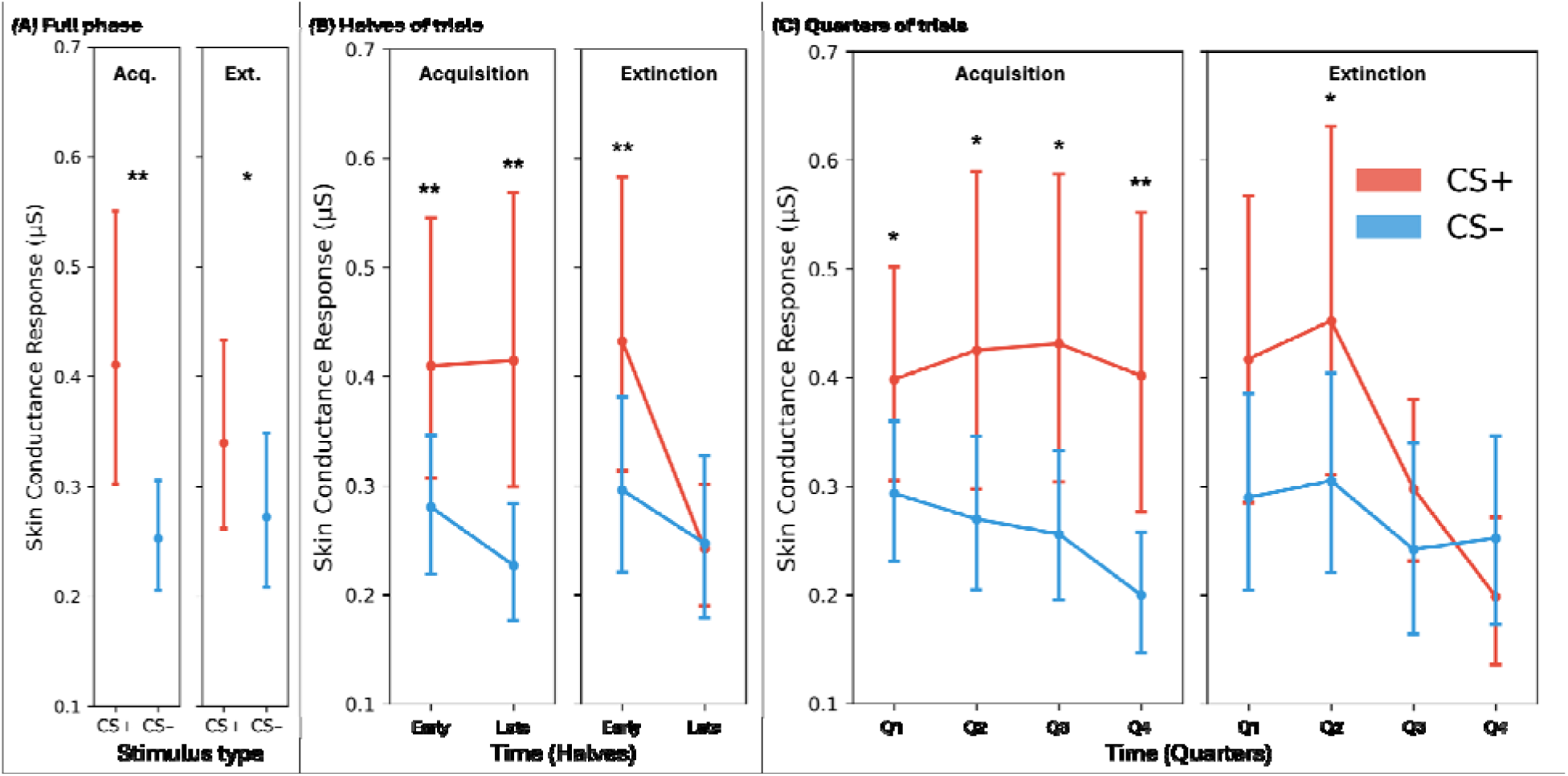
Mean Skin Conductance Responses During Fear Acquisition and Extinction Training. Mean SCRs (in microSiemens, μS) for CS+ and CS– across the full phase (Panel A), halves (Panel B), and quarters (Panel C) for fear acquisition (left sides) and extinction (right sides) training. Error bars represent 95% confidence intervals. Halves were defined as the average of trials 1–8 and 9–16 for fear acquisition training, and 1–4 and 5–8 for extinction training. Quarters were defined as the average of trials 1–4, 5–8, 9–12, and 13–16 for fear acquisition training, and 1–2, 3–4, 5–6, and 7–8 for extinction training. Asterisks denote significant differences between CS+ and CS– for the respective time points (* *p* < .05, ** *p* < .01). SCR = skin conductance response; CS = conditioned stimulus.

#### 3.2.2 Coarse Temporal Dynamics: Early vs. Late Half of Trials

To assess overall conditioned responding across the early and late halves of fear acquisition and extinction training, a two-way repeated-measures ANOVA was conducted to examine the effects of **CS type** (CS+ vs. CS−), **Halves** (early, late), and the **CS type** × **Halves** interaction on SCRs.

During fear acquisition training (See Figure 2B, left panel), the analysis revealed a significant main effect of **CS type**, *F*(1, 113) = 10.50, *p* = .001, partial η*²* = .09, 90% CI [.02, .17]. The main effect of **Halves,** *F*(1, 113) = 1.50, *p* = .22, partial η*²* = .01, 90% CI [.00, .07], and the **CS type × Halves** interaction, *F*(1, 113) = 2.57, *p* = .11, partial η*²* = .02, 90% CI [.00, .08], were not significant. The Bonferroni-corrected post hoc test showed that SCRs were significantly higher for CS+ than for CS– across both early and late halves: **Early CS+**: *M* = 0.40, *SD* = 0.65; **Early CS−**: *M* = 0.28, *SD* = 0.34, 95% CI [.03, .22]; *p* = .008, Cohen’s *d* = 0.25, **Late CS+**: *M* = 0.42, *SD* = 0.74; **Late CS−**: *M* = 0.23, *SD* = 0.29, 95% CI [.08, .30]; *p* = .001, Cohen’s *d* = 0.34.

During extinction training (See Figure 2B, right panel), a similar analysis revealed a significant main effect of **CS type**, *F*(1, 101) = 4.48, *p* = .04, partial η*²* = .04, 90% CI [.01, .12], a significant main effect of **Halves,** *F*(1, 101) = 7.58, *p* = .007, partial η*²* = .07, 90% CI [.01, .16], and a significant **CS type × Halves** interaction *F*(1, 101) = 6.06, *p* = .02, partial η*²* = .06, 90% CI [.01, .14]. The Bonferroni-corrected post hoc tests showed that SCRs were significantly higher for CS+ than CS– only in the early but not the late half: **Early CS+**: *M* = 0.43, *SD* = 0.72; **Early CS−**: *M* = 0.30, *SD* = 0.42, 95% CI [.04, .23]; *p* = .005, Cohen’s *d* = 0.23, **Late CS+**: *M* = 0.25, *SD* = 0.29; **Late CS−**: *M* = 0.25, *SD* = 0.38, 95% CI [–.07, .07]; *p* > 0.99, Cohen’s *d* = −0.01.

#### 3.2.3 Fine-Grained Temporal Evolution: Quarter-Split SCR

Finally, to assess the dynamics of conditioned responding across quarters of trials, a two-way repeated-measures ANOVA was conducted to examine the effects of **CS type** (CS+ vs. CS−), **Quarters** (Q1 through Q4), and the **CS type × Quarters** interaction on SCRs during fear acquisition and extinction training.

During fear acquisition training (See Figure 2C, left panel), the analysis revealed a significant main effect of **CS type**, *F*(1, 112) = 10.52, *p* = .001, partial η*²* = .09, 90% CI [.02, .17]. The main effect of **Quarters,** *F*(3, 336) = 1.66, *p* = .18, partial η*²* = .01, 90% CI [.00, .04], and the **CS type × Quarters** interaction, *F*(3, 336) = 1.36, *p* = .254, partial η*²* = .01, 90% CI [.00, .03], were not significant. The Bonferroni-corrected post hoc tests showed that SCRs were significantly higher for CS+ than CS– across all four quarters: ***Q1* CS+**: *M* = 0.40, *SD* = 0.55; **Q1 CS−**: *M* = 0.30, *SD* = 0.36, 95% CI [.02, .19]; *p* = .03, Cohen’s *d* = 0.23, ***Q2* CS+**: *M* = 0.43, *SD* = 0.81; ***Q2* CS−**: *M* = 0.27, *SD* = 0.37, 95% CI [.03, .28]; *p* = .03, Cohen’s *d* = 0.25, ***Q3* CS+**: *M* = 0.44, *SD* = 0.77; ***Q3* CS−**: *M* = 0.26, *SD* = 0.37, 95% CI [.05, .31]; *p* = .01, Cohen’s *d* = 0.30, ***Q4* CS+**: *M* = 0.40, *SD* = 0.76; ***Q4* CS−**: *M* = 0.20, *SD* = 0.31, 95% CI [.09, .32]; *p* < .01, Cohen’s *d* = 0.34.

During extinction training (See Figure 2C, right panel), the analysis revealed a significant main effect of **CS type**, *F*(1, 100) = 4.48, *p* = .04, partial η*²* = .04, 90% CI [.01, .12], a significant main effect of **Quarters,** *F*(3, 300) = 4.96, *p* < .01, partial η*²* = .05, 90% CI [.01, .08], and a significant **CS type × Quarters** interaction, *F*(3, 300) = 2.75, *p* = .04, partial η*²* = .03, 90% CI [.00, .06]. The Bonferroni-corrected post hoc tests showed that SCRs were significantly higher for CS+ than CS– only in the 2nd but not any other quarter: ***Q1* CS+**: *M* = 0.42, *SD* = 0.75; ***Q1* CS−**: *M* = 0.29, *SD* = 0.47, 95% CI [.00, .26]; *p* = .12, Cohen’s *d* = 0.20, ***Q2* CS+**: *M* = 0.45, *SD* = 0.86; ***Q2* CS−**: *M* = 0.31, *SD* = 0.49, 95% CI [.02, .26]; *p* = .04, Cohen’s *d* = 0.21, ***Q3* CS+**: *M* = 0.29, *SD* = 0.39; ***Q3* CS−**: *M* = 0.24, *SD* = 0.45, 95% CI [−.03, .12]; *p* = .48, Cohen’s *d* = 0.11, ***Q4* CS+**: *M* = 0.19, *SD* = 0.37; ***Q4* CS−**: *M* = 0.25, *SD* = 0.46, 95% CI [−.16, .06]; *p* > .99, Cohen’s *d* = −0.12.

### 3.3 BOLD Activation Across Learning Phases

#### 3.3.1 Conventional Condition-Wise Contrast (All Trials Averaged)

To establish a solid comparability of our novel trial segments fMRI analysis at 0s, 2s, and 4s relative to the CS onset, we analyzed the fMRI averaged across all trials of fear acquisition and extinction training, prior to splitting trials into early and late halves (described in section 3.3.2 below) and quarters of trials Q1-Q4 (described in section 3.3.3 below).

##### Fear Acquisition Training

###### CS+ > CS– Contrast

The CS+ > CS– differential response averaged across all acquisition trials demonstrated a dynamic temporal progression. At CS onset (0s), activation was prominent in regions associated with salience and visual processing, including the dACC, anterior and posterior callosal body, fornix, visual cortex (V4, V5), insular cortex (predominantly left), and bilateral operculum (Supplementary Figure S1, top). By 2s, the response shifted toward a broader lateralized network encompassing the bilateral visual cortex (V4), left parietal and frontal operculum, left Broca’s area, left precentral gyrus, left secondary somatosensory cortex, left frontal pole, and right frontal operculum (Supplementary Figure S1, middle). At 4s, the threat response became highly localized, restricted entirely to the left frontal operculum and left frontal pole (Supplementary Figure S1, bottom).

###### CS– > CS+ Contrast

The CS– > CS+ contrast revealed an expanding, widespread network representing the safety signal. At 0s (Supplementary Figure S2, top), initial activations were observed in the left parahippocampal gyrus, bilateral visual cortex (V1-V4), right inferior parietal lobule, and left superior parietal lobule. By 2s (Supplementary Figure S2, middle), this response broadened into major global activations characterized by robust engagement of the bilateral vmPFC, bilateral cuneus, left hippocampus and parahippocampus, bilateral operculum, and bilateral inferior parietal lobule. At 4s (Supplementary Figure S2, bottom), extensive activation persisted, shifting notably to motor and regulatory regions, including the dACC, bilateral primary motor cortex, bilateral premotor cortex, and bilateral vmPFC.

##### Extinction Training

###### CS+ > CS– Contrast

Throughout extinction training, the CS+ > CS– contrast yielded no significant differential activations across the 0s and 2s temporal segments (Supplementary Figure S3, top and middle), which might reflect a successful attenuation of the fear response, even occurring within the 8 trials implemented. At 4s, however, a significant activation in bilateral frontal operculum and insula has been found (Supplementary Figure S3, bottom).

###### CS– > CS+ Contrast

The contrast CS– > CS+ during extinction training revealed a sustained and progressive engagement of visual and associative processing regions. At 0s, activation was narrowly localized to the bilateral visual cortex (V1-V4) and right dmPFC as part of the superior frontal gyrus (Supplementary Figure S3, top). By 2s, the response expanded to encompass a broader activation in the bilateral visual cortex (V1-V4) and bilateral dmPFC, bilateral hippocampus and parahippocampus, as well as bilateral cuneal and precuneal cortices (Supplementary Figure S3, middle). At 4s, activation progressed further along the ventral stream, recruiting the bilateral temporal and occipital cortices, with bilateral visual cortex V1-V4, bilateral cuneus and precuneus cortices, Left amygdala, bilateral hippocampus (stronger left activation), right primary somatosensory and motor cortices, bilateral dmPFC and vmPFC (Supplementary Figure S3, bottom)

#### 3.3.2 Coarse Temporal Dynamics: Early vs. Late Trials

To examine the spatiotemporal dynamics of fear learning at a broader temporal scale, we analyzed BOLD responses at three time points (0s, 2s, 4s relative to CS onset) separately for the first and second half of trials (Early: trials 1-8; Late: trials 9-16 for fear acquisition training; Early: trials 1-4; Late: trials 5-8 for extinction training).

##### Fear Acquisition Training

###### CS+ > CS– Contrast

The CS+ > CS– differential response revealed a progression of network engagement from early to late acquisition. During early acquisition, widespread salience and sensory activations were observed at CS onset (0s), encompassing the dACC, PCC, fornix, bilateral thalamus, bilateral secondary somatosensory cortex, and operculum (central and parietal; Supplementary Figure S1, top). By 2s (Supplementary Figure S1, middle), activation shifted to the right central/frontal operculum and insula, bilateral visual cortex (V1-V4), right ventrolateral prefrontal cortex (vlPFC), and right primary somatosensory cortex extending to the inferior-anterior parietal lobule. At 4s (Supplementary Figure S1, bottom), the response was isolated to the right frontal operculum, insula, and right vlPFC.

During late acquisition, the network demonstrated reduced and more focused engagement. At 0s (Supplementary Figure S1, top), activation was limited to the dACC and left operculum. By 2s (Supplementary Figure S1, middle), activation persisted in the right vlPFC, right insula, and frontal operculum. At 4s (Supplementary Figure S1, bottom), only the right vlPFC remained active, reflecting a refinement of the fear response into sustained executive maintenance over time.

###### CS– > CS+ Contrast

The CS– > CS+ contrast revealed an expanding pattern of activation for the safety signal as acquisition progressed. During early acquisition, 0s activation was limited to the right inferior parietal lobule (Supplementary Figure S2, top). At 2s (Supplementary Figure S2, middle), this expanded into a broad network including the right inferior parietal lobule, bilateral visual cortex (V1-V4), bilateral premotor cortex, cingulate gyrus, primary motor cortex, left insula, operculum, primary somatosensory cortex, vmPFC, and vlPFC. By 4s (Supplementary Figure S2, bottom), activation encompassed the bilateral premotor cortex, primary/secondary somatosensory cortex, left inferior parietal lobule, left V5, bilateral vmPFC, left hippocampus, thalamus, and fornix.

During late acquisition, the safety signal recruited extensive emotion regulation and memory networks. At 0s (Supplementary Figure S2, top), activation involved the left hippocampus, parahippocampus, visual cortex (V2-V4), cuneus, precuneus, right inferior parietal lobule, insula, and operculum. By 2s (Supplementary Figure S2, middle), this response peaked with massive engagement of the bilateral hippocampus, parahippocampus, amygdala, visual cortex (V1-V4), cuneus, precuneus, thalamus, brainstem, vmPFC, premotor cortex, operculum, insula, and cerebellar vermis and crus. At 4s (Supplementary Figure S2, bottom), sustained activation was observed in the bilateral premotor cortex, primary/secondary somatosensory cortex, left inferior parietal lobule, left V5, bilateral vmPFC, left hippocampus, thalamus, fornix, and amygdala.

##### Extinction Training

###### CS+ > CS– Contrast

During early extinction, no significant differential activation emerged at 0s or 2s (Supplementary Figure S3 top and middle). However, at 4s, significant activation was observed in the bilateral frontal operculum and insula (Supplementary Figure S3 bottom). By late extinction, the differential response was completely habituated, with no significant activation observed at 0s, 2s, or 4s. However, this lack of significant activation is likely a consequence of averaging out transient signals across the entire phase; as detailed in our subsequent quarters analysis (Section 3.3.3), this global average obscures a distinct, temporary resurgence of fear network activation during the second quarter (Q2) of extinction as reflected in both SCRs and fMRI results.

###### CS– > CS+ Contrast

The CS– > CS+ contrast during extinction training showed early, widespread engagement of memory and visual networks that diminished over time. During early extinction, 0s activation involved the bilateral visual cortex (V1-V4), right cerebellar crus I and II, vermis VI, and left middle frontal gyrus (Supplementary Figure S4, top). At 2s, this expanded to include extensive bilateral visual cortex, cuneus, precuneus, left amygdala, bilateral hippocampus, and vmPFC (Supplementary Figure S4, middle). By 4s, activation encompassed the bilateral hippocampus, parahippocampus, left amygdala, bilateral cuneus, precuneus, vmPFC, dmPFC, bilateral visual cortex (V1-V5), and cerebellum (Supplementary Figure S4, bottom).

During late extinction, the network response became more restricted. At 0s, activation was limited to the bilateral visual cortex (V1-V4; Supplementary Figure S4 top). By 2s, this included the bilateral visual cortex (V1-V4), bilateral cuneus, left precuneus, and left hippocampus (Supplementary Figure S4, middle). At 4s, sustained activation was observed in the bilateral cuneus, right precuneus, superior parietal lobule, and cerebellum (Supplementary Figure S4, bottom).

#### 3.3.3 Fine-Grained Temporal Evolution: Quarter-Split Model

To examine the spatiotemporal dynamics of fear learning with higher temporal resolution, we analyzed BOLD responses at three time points (0s, 2s, 4s relative to CS onset) separately for each quarter of trials (Q1: trials 1-4; Q2: trials 5-8; Q3: trials 9-12; Q4: trials 13-16 for fear acquisition training; Q1: trials 1-2; Q2: trials 3-4; Q3: trials 5-6; Q4: trials 7-8 for extinction training). We compared the results of our quarter-split analysis to the combined statistical maps from Fullana et al. (2016) and Wen et al. (2024), matched by contrast direction (e.g. CS+ > CS–) and trial blocks (Q1-Q4), reported in Supplementary Figures S5-S12. The green contours around significant clusters of voxels in Supplementary Figures S5, S7, S9 and S11 delineate a spatial overlap of the significant brain activations from our analysis with the combined Fullana et al., (2016) and Wen et al., (2024) statistical maps. The blue contours in Supplementary Figures S6, S10 (CS+ > CS– contrasts), and the red contours in Supplementary Figures S8, S12 (CS– > CS+ contrasts) are equivalently delineating the spatial boundaries of the significant regions uniquely present in our analysis and not in the combined meta-analytic reference maps (Fullana and Wen). The black contours in Figures S6, S8, S10, and S12 delineate the spatial boundaries of the significant regions uniquely present in our analysis and not in the combined meta-analytic reference maps (Fullana and Wen).

##### Fear Acquisition training

###### CS+ > CS– contrast

The CS+ > CS– differential response showed a three-phase temporal evolution that shifted across trial quarters (Figure 3, red activations, Figure S1, and Table S3). During Q1 acquisition, activation at CS onset (0s) was restricted to the right visual cortex (V4/V5). This was followed by the recruitment of the right frontal operculum, right vlPFC, and sustained visual cortex activation at 2s, resolving into sustained right vlPFC activation by 4s. Analysis of Q2 revealed maximal network engagement. At 0s, extensive limbic-striatal activation was observed, encompassing the dACC, bilateral hippocampus, parietal operculum, striatum, and cerebellum. By 2s, this activation transitioned to a frontoparietal control network (right dlPFC, insula, and bilateral inferior parietal lobule), ending with sustained activation of the right vlPFC and frontal operculum at 4s. During Q3, the network refined further, showing dACC and somatosensory-motor engagement at 0s and frontoparietal activation (supramarginal gyrus, dlPFC) at 2s, before isolating to the right vlPFC at 4s. By late acquisition (Q4), no significant differential activation emerged at 0s or 2s, with only the right vlPFC remaining active at 4s.

**Figure 3.**
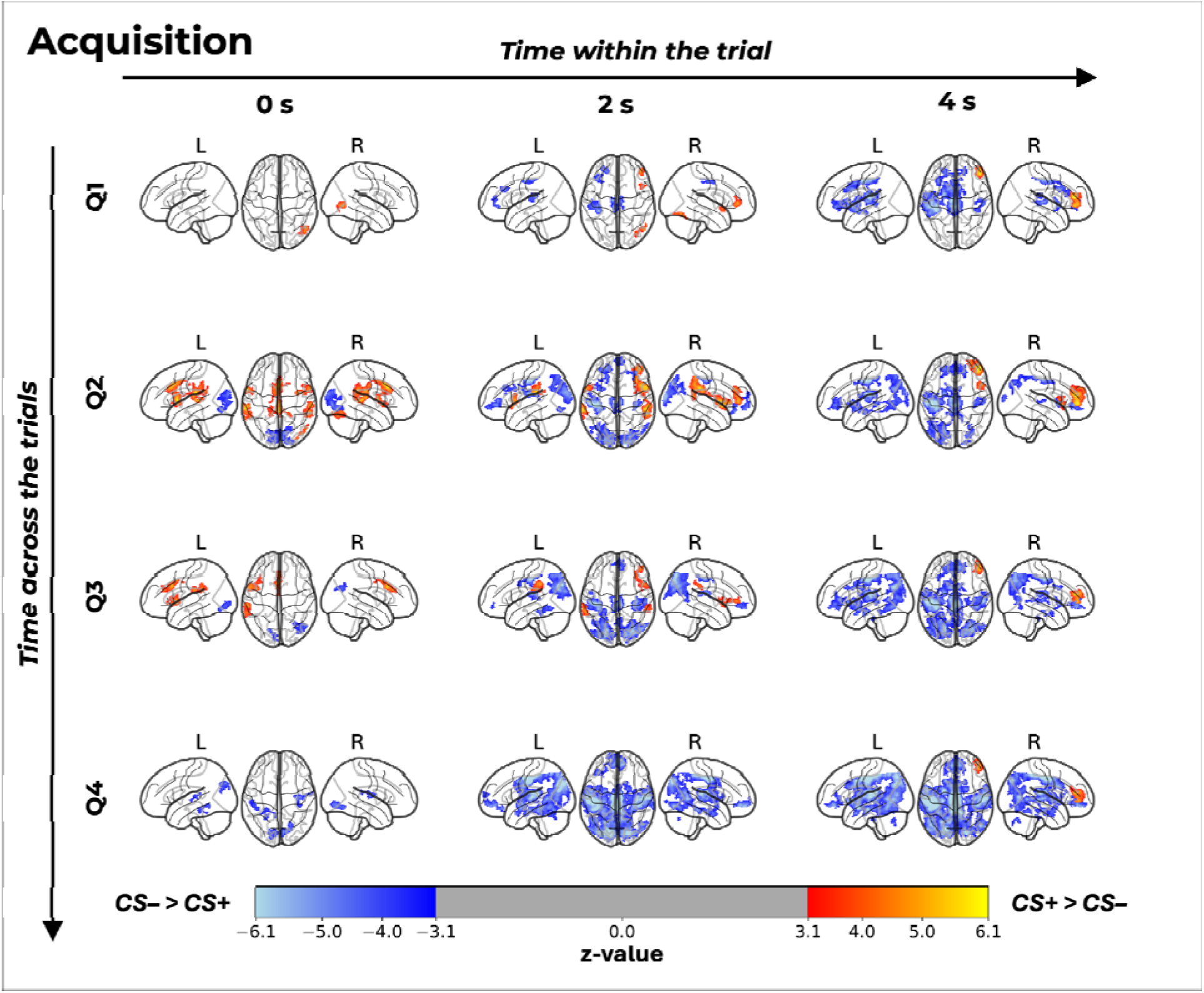
Spatiotemporal dynamics of BOLD activation during fear acquisition training. The grid illustrates the evolution of neural responses for the CS+ > CS– contrast (represented in red/yellow) and the CS– > CS+ contrast (represented in blue) during fear acquisition training. The data is organized by acquisition quarters (rows Q1–Q4) to show changes across the trials, and modeled at three latencies post-stimulus onset (columns: 0 s, 2 s, and 4 s) within the trial. “Not Significant” indicates the absence of suprathreshold clusters in our data for that specific time point and quarter. All activation maps are thresholded and plotted on a transparent MNI152 template showing lateral (L/R) and superior views. The colorbar indicates z-statistic values ranging from 3.1 to 6.1 for CS+ > CS– and from −6.1 to −3.1 for the CS– > CS+.

While our 0s segment in Q1 shows spatial overlap with the combined meta-analytic maps from Fullan and Wen (Supplementary Figure S5), this overlap diverges at the 2s and 4s segments. Most notably, the late-latency right vlPFC engagement observed in our data is entirely absent from the established reference maps (Supplementary Figure S6)

###### CS– > CS+ contrast

The response to the safety signal (CS–) exhibited a delayed onset and progressive recruitment of safety-signaling regions over time (Figure 3, blue activations, Figure S2, and Table S3). In the earliest phase (Q1), no significant differential activation was observed at 0s or 2s. However, by Q2 and Q3, differential activation emerged in the visual processing stream (V1–V4, cuneal cortex) and right inferior parietal lobule at 0s and 2s, suggesting enhanced visual attention to the safety cue. By the final quarter (Q4), the CS– evoked a robust safety signaling pattern at 0s and 2s, recruiting the left hippocampus, parahippocampal gyrus, and precuneal/cuneal cortices, regions consistent with contextual safety encoding and Default Mode Network (DMN) engagement. Notably, at the late latency time point (4s), a widespread activation pattern was observed across all quarters (Q1–Q4). This late-phase response overlapped primarily with the canonical fear network described by Fullana et al. (2016), suggesting a generalized salience or rebound response to the safety cue at the end of the trial window.

##### Extinction training

###### CS+ > CS– contrast

In contrast to fear acquisition training, the CS+ elicited minimal differential activation during the Q1, Q3, and final Q4 phases of extinction training, with no significant clusters observed at any time point (0s, 2s, 4s; Figure 4, red activations, Figure S3 and Table S4). However, a distinct, transient resurgence of fear network activation emerged during Q2. While the onset (0s) remained quiescent, the 2s time point revealed robust activation in the salience network, including the bilateral insula (predominantly left), bilateral operculum, and left thalamus extending to the right caudate, alongside the vlPFC. This engagement persisted into the late processing window (4s), where activation remained anchored in the bilateral frontal operculum, left insula, and vlPFC, indicative of a temporary reinstatement of fear processing before subsequent suppression in Q3 and Q4.

**Figure 4.**
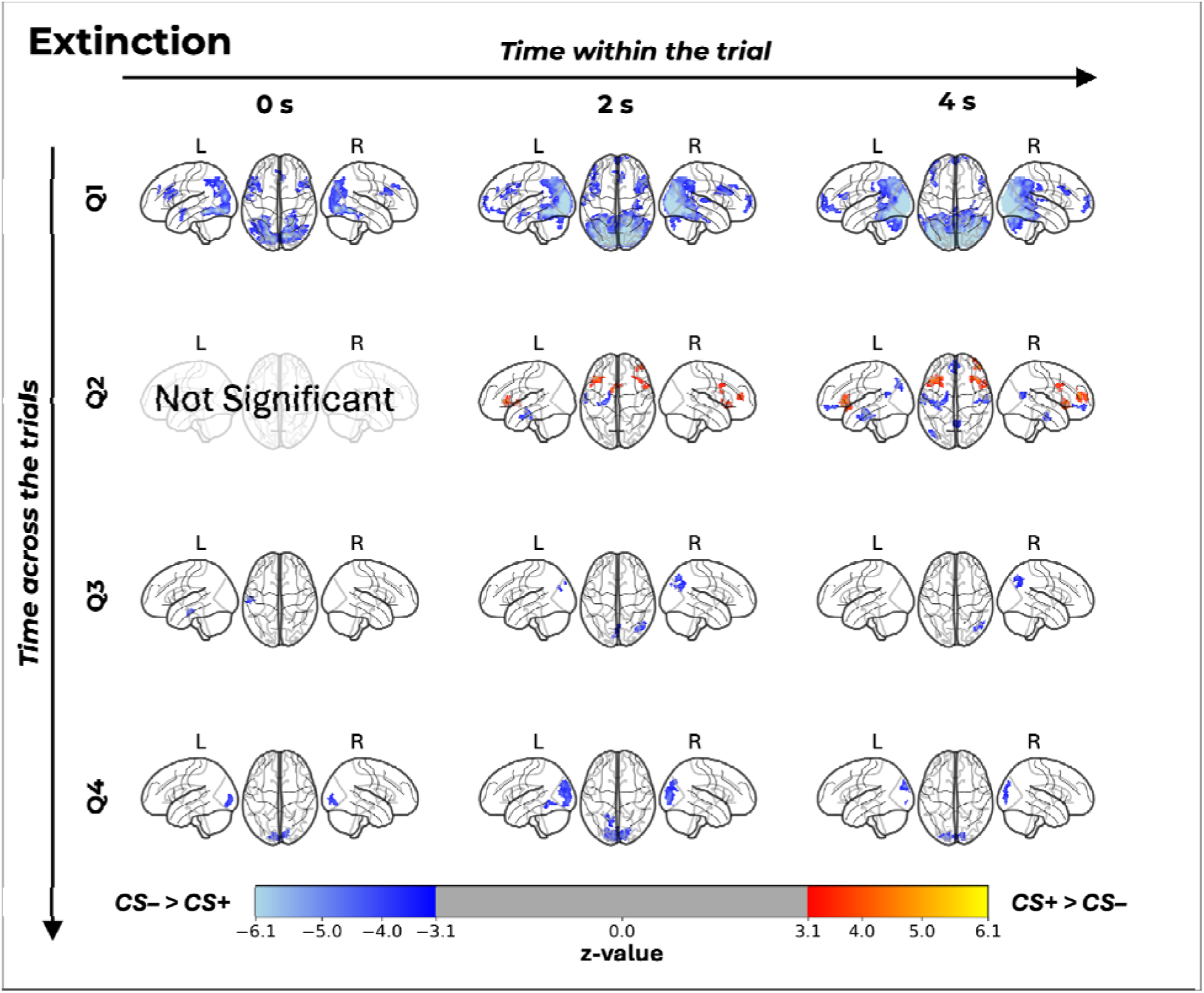
Spatiotemporal dynamics of BOLD activation during extinction training. The grid illustrates the evolution of neural responses for the CS+ > CS– contrast (represented in red/yellow) and the CS– > CS+ contrast (represented in blue) during extinction training. The data is organized by extinction quarters (rows Q1–Q4) to show changes across the trials, and modeled at three latencies post-stimulus onset (columns: 0 s, 2 s, and 4 s) within the trial. “Not Significant” indicates the absence of suprathreshold clusters in our data for that specific time point and quarter. All activation maps are thresholded and plotted on a transparent MNI152 template showing lateral (L/R) and superior views. The colorbar indicates z-statistic values ranging from 3.1 to 6.1 for CS+ > CS– and from −6.1 to −3.1 for the CS– > CS+.

###### CS– > CS+ contrast

In Q1, the response was robust across all time points. At onset (0s), activation encompassed the posterior parahippocampus, paracingulate cortex, and bilateral cuneal/precuneal cortices (Figure 4, blue activations, Figure S4 and Table S4). This evolved at 2s and 4s to include the vmPFC (region critical for extinction memory consolidation) alongside the bilateral hippocampus, posterior cingulate cortex, and extensive cerebellar engagement (Crus I/II). During Q2, the temporal dynamics shifted. While processing at 0s yielded no significant differences, the late window (4s) revealed a unique integration of emotional and memory circuits. Specifically, the CS– recruited the left amygdala, bilateral hippocampus, and vmPFC, alongside the left inferior parietal lobule. By the latter quarters of extinction (Q3 and Q4), differential activation transitioned toward sensory and attentional processing. Q3 was characterized by parietal engagement (right superior and inferior parietal lobules) at 2s and 4s. Finally, Q4 showed activation restricted almost exclusively to the visual cortex (V1–V4) and cuneus across all time points, reflecting habituation of higher-order safety processing mechanisms.

## 4. Discussion

In the present study, we examined the spatiotemporal dynamics of fear acquisition and extinction learning using a temporally resolved fMRI analysis. By modeling BOLD responses at three within-trial latencies (0 s, 2 s, and 4 s relative to CS onset) and across four quarters of trials, we aimed to reveal transient neural activations that are obscured by conventional trial-averaged or phase-averaged analyses. In line with this approach, we evaluated conditioned responses at three temporal scales, averaged across the full phase, split into early and late halves, and divided into individual quarters (Q1–Q4), for both SCRs and fMRI BOLD signals. The following sections discuss these findings in the context of established fear and safety networks, highlight novel activation patterns not captured by prior meta-analyses, and address the clinical and methodological implications of a temporally resolved approach to fear learning.

### Contingency ratings and SCRs

Self-report questionnaire results confirm that participants successfully acquired and updated explicit contingency knowledge. After fear acquisition training, estimates of CS+ reinforcement (∼58%) closely mirrored the actual 62.5% schedule, demonstrating a clear cognitive differentiation between fear (CS+) and safety (CS−) signals. Following extinction training, participants unanimously reported zero stimulations, indicating full conscious awareness that the US had been removed. While these ratings verify explicit declarative memory of the contingencies, we analyzed SCRs to assess corresponding changes in implicit physiological arousal during the phases. Because autonomic responses change dynamically across training, we evaluated SCRs across three temporal resolutions: averaged across all trials of acquisition and extinction training, split into early and late halves of respective training phases, and divided into quarters (Q1–Q4). Averaging across all trials initially suggested a strong, persistent differential effect (CS+ > CS−) throughout both fear acquisition and extinction training (see Figure 2A). However, finer temporal subdivisions revealed a much more localized progression. Splitting the data into early and late halves showed the differential response persisted throughout fear acquisition training but was restricted to early extinction, with CS+ and CS– responses equalizing by late extinction (see Figure 2B). The quarter-split analysis further refined this trajectory (see Figure 2C). The CS+ > CS– effect remained consistent across all acquisition quarters (reaching its highest significance in Q4), whereas during extinction training, the differential response was only significant in Q2. Notably, we also observed an elevated response to the CS– in Q2. As discussed in our previous work using this identical paradigm (Gabdulkhakov et al., 2026), this elevation likely reflects ambiguity introduced by the context switch from fear acquisition to extinction training. Consequently, the apparent persistence of differential fear across all extinction trials is actually an artifact of a strong, localized effect in Q2 that has smeared across the broader averaging window.

### Temporal Evolution of Fear Acquisition: From Sensory Salience to Regulation

Our quarter-split analysis revealed a distinct three-phase progression of neural engagement during fear acquisition training (Figure 3 and Table S3), challenging static models of the “fear network” (Fullana et al., 2016, 2018; Wen et al., 2024). Phase 1 (Sensory Dominance, Q1): Early learning was characterized by activation restricted primarily to the visual cortex (V4/V5) at CS onset. This suggests that the initial stage of fear learning is driven by sensory feature extraction and heightened perceptual salience of the CS+ before robust affective associations are fully consolidated (Miskovic & Keil, 2012). Phase 2 (Associative Peak, Q2): The second quarter marked the peak of associative learning, recruiting a widespread network including the dACC, bilateral hippocampus, striatum, and insula. This “explosion” of activation likely reflects the maximal encoding of the prediction error and the formation of the CS/US contingency. Phase 3 (Refinement and Maintenance, Q3–Q4): As training progressed, the broad limbic-striatal response rapidly habituated. By Q4, no significant activation was observed at trial onset (0s). Instead, the response was characterized by sustained activation of the right vlPFC at the 4s time point. This shift from limbic reactivity to prefrontal engagement suggests a transition from active threat detection to the maintenance of learned rules and top-down regulation.

### Within-Trial Dynamics: The Shift from Detection to Regulation

The higher temporal resolution of our analysis (0s, 2s, and 4s relative to CS onset) uncovers a dynamic functional shift within individual trial segments. Across fear acquisition training, the CS+ response consistently began with sensory-motor engagement (0s) and transitioned to frontoparietal control networks (2s). Crucially, the late-latency window (4s) in fear acquisition training was defined by sustained right vlPFC activation across all quarters. Unlike the transient dACC or insula activation associated with immediate threat appraisal, this persistent vlPFC engagement suggests ongoing top-down emotion regulation (Cheng et al., 2022; Ochsner et al., 2012; Wager et al., 2007) and/or inhibitory control and action readiness (Aron et al., 2014; Swick et al., 2014; Vinberg et al., 2022). Historically, rodent models have primarily implicated the vlPFC in extinction recall, rather than during fear acquisition training (Tian et al., 2011). However, our ability to isolate delayed temporal segments reveals that right vlPFC activation likely represents the active maintenance of threat prediction or the preparatory inhibition of motor responses prior to expected US delivery. Notably, we observed a similar 4s-latency vlPFC activation during the second quarter (Q2) of extinction. This neural signature temporally maps onto the transient return of differential SCRs observed in extinction Q2, reinforcing the role of the vlPFC in mobilizing regulatory control during periods of heightened autonomic arousal.

### Active Safety Learning and the “Relief” Response (CS– > CS+)

A striking finding in our data is the emergence of a robust CS– > CS+ profile towards the end of the trial in early quarters (i.e., 4s and Q1) and towards the final quarter of fear acquisition training (Q4). Rather than being treated as a neutral baseline (Rescorla, 1967; Rescorla, 1988), the CS– elicited strong activation in the hippocampus, parahippocampal gyrus, and precuneus. This supports the safety signal hypothesis, positing that the CS– acquires explicit inhibitory value (Christianson et al., 2012). The recruitment of the hippocampus specifically points to the contextual encoding of safety-learning (Labrenz et al., 2022), where and when the threat will not occur. Furthermore, at the late-trial window (4s), the CS– elicited widespread activation overlapping with the canonical salience network (Fullana et al., 2016). We interpret this paradoxical finding as a “relief” or “omission” response (Dunsmoor et al., 2012). When the expected timing of the US passes without an electrical stimulation, the massive prediction error (unexpected safety) generates a salience signal that anatomically mimics threat processing but functionally represents relief. Interestingly, the ‘relief’ response to the safety cue was also characterized by robust engagement of primary and secondary somatosensory cortices, as well as the premotor cortex. This suggests that the omission of the anticipated electrical stimulus prompts a tactile re-evaluation and motor relaxation, actively mapping the absence of threat onto the body schema (Wiemer et al., 2024).

### Extinction Dynamics: Spontaneous Recovery and the “Extinction Burst”

The transition to extinction training was marked by the successful elimination of the CS+ threat response in Q1, Q3, and Q4. However, our data reveal a significant non-linear phenomenon in Q2, characterized by a transient resurgence of CS+ activation in the bilateral insula, frontal operculum, and dlPFC (2s and 4s). This pattern is also supported by the return of differential CS+ > CS– effect in SCRs, which shows a direct link between the two modalities under closer, time-resolved investigation. Thus, extinction learning seems to follow a monotonic decay of fear but to involve a mid-phase period of cognitive conflict or prediction error re-evaluation. The specific involvement of the insula and dlPFC during this burst points to a re-engagement of salience and executive control networks to resolve the ambiguity of the no-shock outcome after the initial “easy” extinction in Q1.

### Mechanisms of Safety Consolidation During Extinction

While the CS+ response diminished, the CS– recruited a robust safety-signaling network, particularly involving the vmPFC and hippocampus during the windows at 4s of Q1 and Q2. The engagement of the vmPFC is critical, as this region is the canonical locus of extinction retention and down-regulation of the amygdala. Interestingly, in Q2 (4s), we observed simultaneous activation of the left amygdala and vmPFC for the CS– > CS+ contrast. While amygdala activation is typically associated with threat, its recruitment here (alongside the vmPFC) likely reflects the active processing of positive safety salience or the consolidation of the “safe” memory trace. By Q3 and Q4, this active processing habituated, leaving only sensory (visual) activation, indicating that the safety memory had been fully consolidated and no longer required top-down resources. Alongside prefrontal-limbic engagement, we observed robust cerebellar activation (specifically Crus I and II) during the CS− presentation in early extinction. Recent predictive coding frameworks suggest the cerebellum acts as a critical comparator, generating prediction errors when expected aversive outcomes are omitted (Doubliez et al., 2023; Ernst et al., 2019). Its co-activation with the vmPFC highlights a distributed network for computing and consolidating the safety memory trace.

### Dendrograms of fMRI BOLD activations: from whole trials to quarters

The SCR results in Figure 2A-C, compared to dendrograms in Figures S1-S4, show the core dynamics of learning captured in both imaging modalities. Most prominently, as mentioned above, the corresponding differential effect in Q2 of extinction for SCRs (Figure 2) is mirrored by the fMRI results (Figure 4 and Supplementary Figure S3). Specifically, we observed corresponding activations in the bilateral frontal operculum and insula during the 2s trial segment of Q2, as well as in the 4s segment during both the early half and Q2 of the phase. Furthermore, examining the temporal progression of the CS+ > CS– contrast (Supplementary Figure S1) suggests that the 2s segment represents a transitional state. It encompasses a functional overlap of early (0s) and late (4s) activations. Notably, the classical areas such as dACC remain consistently specific for the 0s segment, which might point to the limitation of focusing solely on the response towards the CS at 0s post-onset, as the engagement of the areas we reported above at 2s and 4s might be systematically overlooked.

We compared our results to the combined statistical maps from Fullana et al. (2016) and Wen et al. (2024), matched by contrast direction (CS+ > CS–) and trial blocks. While our 0s segment in Q1 shows spatial overlap with the combined meta-analytic maps (Supplementary Figure S5), this overlap diverges at the 2s and 4s segments. Most notably, the late-latency right vlPFC engagement observed in our data is entirely absent from the established reference maps (Supplementary Figure S6). This divergence is expected, as the combined reference maps derived from analyses strictly locked to the CS onset. Comparing the unique activations to established meta-analyses highlights the limitations of static full-trial average or all-trial-average contrasts. For example, our segmented approach isolates transient inhibitory control nodes, such as the right vlPFC at the 4s latency, which are completely averaged out in the combined Fullana and Wen maps.

As illustrated in Supplementary Figures S5 through S12, the combined reference maps from Fullana and Wen capture a macroscopic, brain-wide network of fear and safety processing, largely because they reflect standard, single-time-point or trial-averaged analyses. In contrast, our temporally segmented approach provides a high-resolution temporal lens, revealing that these broad regional activations are actually driven by highly specific, time-dependent neural cascades. For instance, while our 0s segment strongly overlaps with the established literature, our later trial segments successfully isolate precise, transient nodes - such as the late right vlPFC engagement - that are otherwise entirely averaged out in broader meta-analytic maps. These newly identified areas should be replicated in future studies. If confirmed, they could represent important targets for modulation using brain stimulation techniques, with the aim of influencing the learning process and, ultimately, informing interventions for the treatment of anxiety disorders.

The robust, diagonal spatiotemporal activation pattern observed for the CS− > CS+ contrast (Figure S2) reflects the progressive neural differentiation of safety and threat across both the fear acquisition training and the individual trial timeframe. Early in fear acquisition training, the absence of significant differentiation aligns with the initial ambiguity of stimulus contingencies before associative learning has consolidated. As learning progresses towards the final blocks, we observe a broad cortical recruitment primarily mapping onto the Default Mode Network (DMN), including regions such as the vmPFC and hippocampus, which are established nodes for safety signaling and threat inhibition (Marstaller et al., 2016). This across-trial evolution is compounded by the within-trial unfolding of the BOLD signal, where the maximal activation observed at 4 seconds captures the delayed hemodynamic manifestation of cognitive threat appraisal. Together, these dynamics suggest that the pronounced CS− > CS+ maps in the latter stages of fear acquisition training are driven both by the active neural evaluation of the safety cue and the concurrent suppression of the DMN by the salient, aversive CS+.

### Limitations

One of the main limitations of this study is that, despite focusing on trial segments based on the literature (Gabdulkhakov et al., 2026; Starita et al., 2022), these trial segments separated by 2s are arbitrary and might not have strict relevance to the millisecond-level timing of neurophysiological processes. We strongly believe that future neuroimaging studies utilizing faster sequences (e.g., Feinberg et al., 2010), combined with source-localized or simultaneously recorded EEG, will reveal even finer temporally specific activation of these areas. Conversely, the trial segment at 2s relative to CS onset may be temporally too close to the 0s and 4s segments, resulting in overlapping activations from both. Notably, however, this trial segment at 2s relative to the CS onset lacks a classical activation in fear-related regions like the dACC, present at 0s trial segment. The restriction of dACC engagement strictly to the 0s segment clarifies why this region so heavily dominates prior literature, which has traditionally relied on CS onset-locked or trial-averaged analyses. Another possible limitation is that more recent studies, such as Wen et al. (2024), have employed MVPA, whereas the present study relies on univariate analyses. Applying an MVPA approach to the trial segments in future work could therefore reveal whether the underlying neural dynamics follow a different pattern, though such an analysis falls beyond the scope of the present manuscript.

## 5. Conclusion

In conclusion, we demonstrated the critical importance of analyzing distinct trial segments and phases of learning in fMRI fear conditioning research. A central motivation for this approach was the puzzling absence of amygdala and vmPFC activation in the Fullana et al. (2016) meta-analysis, regions canonically implicated in fear and safety learning of animal literature. By tracing the BOLD signal dynamically in segments within and across trials, we show that these activations were not absent but temporally displaced: amygdala and vmPFC responses emerged selectively in the 2s and 4s segments following CS onset, and would have been diluted or missed entirely by analyses locked to CS onset or averaged across the full trial window. This finding directly extends the high-resolution block analysis of Wen et al. (2024), to whom we add a quarterly split of extinction training and a delineation of the spatial evolution of the safety (CS− > CS+) contrast within trial segments and across quarters of trials. Taken together, our results argue that the “fear network” is not a monolithic, static entity, but rather a fluid cascade of sensory detection, associative peak firing, and prefrontal maintenance that shifts dramatically from the first to the final trial, and critically, across different temporal proximities relative to the CS onset.

## Author contribution

**Arslan Gabdulkhakov**: Methodology; Software; Formal analysis; Data curation; Visualization; Writing – original draft; Writing – review & editing.

**Christian J. Merz**: Methodology; Investigation; Formal analysis; Writing – review & editing.

**Christoph Fraenz**: Investigation; Data curation.

**Erhan Genç**: Conceptualization; Methodology; Project administration; Supervision; Funding acquisition; Data curation; Writing – review & editing.

## Acknowledgements

We would like to thank our student assistants, Patrick Friedrich and Helene Selpien, for supporting data recordings. We also would like to thank Tobias Otto and Dorothea Metzen for assistance with the analysis of SCR data.

## Data Availability Statement

The preprocessed fMRI statistical maps are available at https://osf.io/jn7b8. Raw data and other materials are available upon request.

## Use of AI-generated content (AIGC) and tools

We used Perplexity Pro to improve grammar, enhance clarity, and check for cohesion at the sentence level in the manuscript.

