## Supplementary Materials for "BOLD signals of learning dynamics across and within trials beyond the classical fear and extinction network"

**for**

Prof. Dr. Erhan Genç

Neuroimaging and Interindividual Differences Unit

Department of Psychology and Neurosciences

Leibniz Research Centre for Working Environment and Human Factors (IfADo)

Ardeystraße, 67

44139 Dortmund, Germany

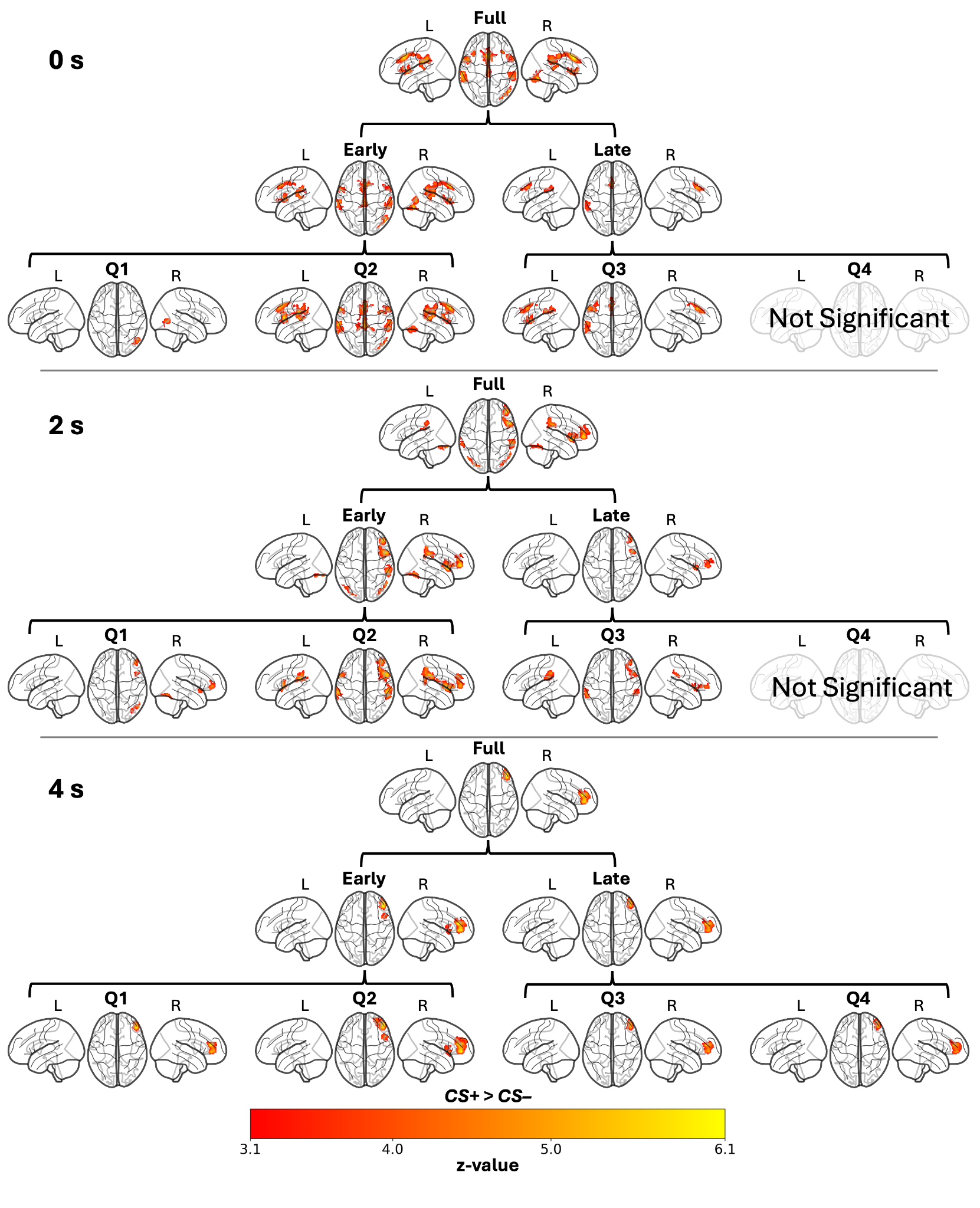

*Figure S1*. Hierarchical BOLD activation patterns for the CS+ > CS– contrast during fear acquisition training, modeled at 0 s, 2 s, and 4 s post-stimulus latencies. The dendrogram structure illustrates the decomposition of the analysis from the Full Training phase (top in each panel), to Halves (middle in each panel), and finally to Quarters (bottom in each panel; Q1–Q4). "Not Significant" indicates the absence of suprathreshold clusters. All maps are plotted on a transparent MNI152 template. The red to yellow color bars code the thresholded z-statistic value from 3.1 to 6.1.

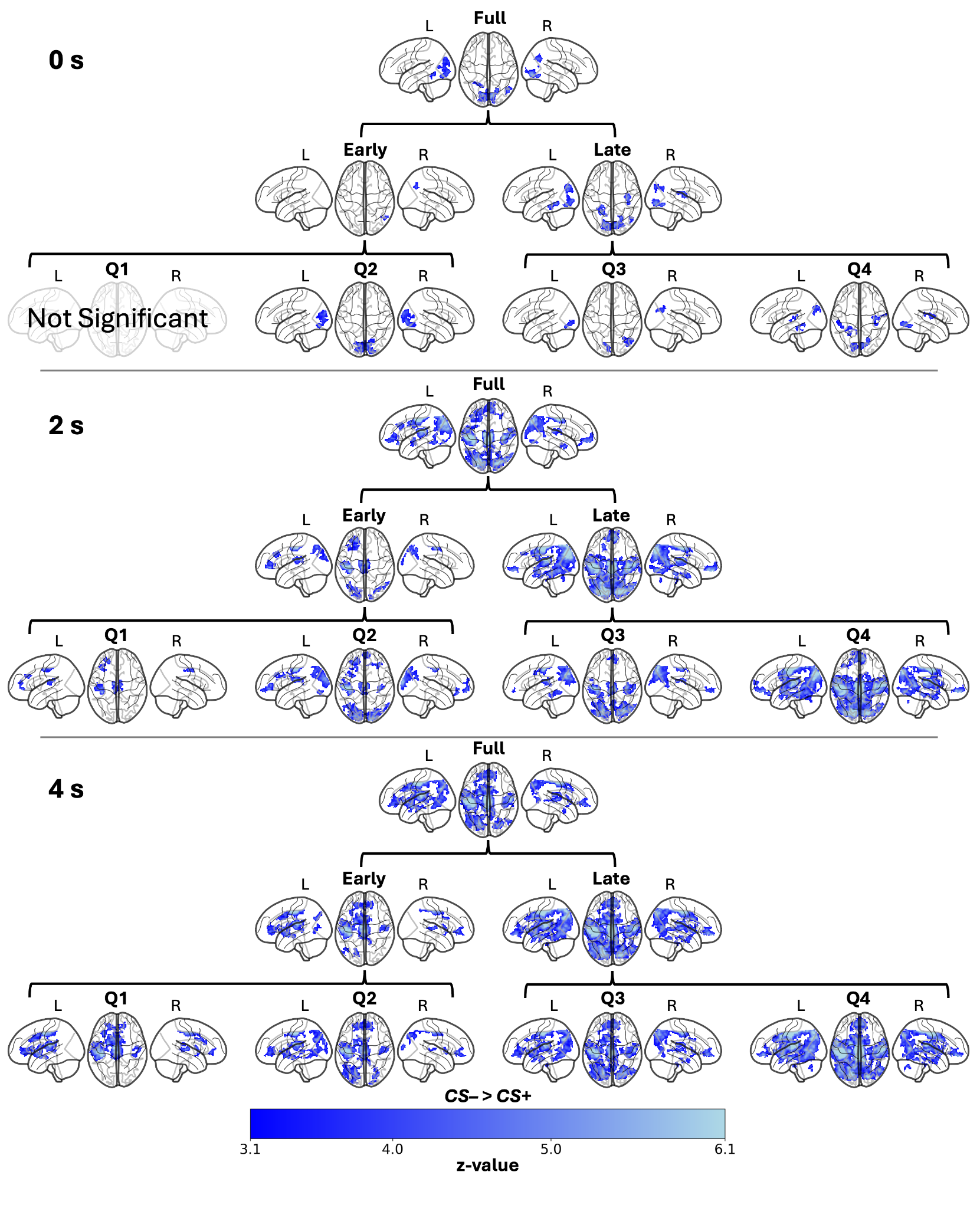

*Figure S2*. Hierarchical BOLD activation patterns for the CS– > CS+ contrast during fear acquisition training, modeled at 0 s, 2 s, and 4 s post-stimulus latencies. The dendrogram structure illustrates the decomposition of the analysis from the Full Training phase (top in each panel), to Halves (middle in each panel), and finally to Quarters (bottom in each panel; Q1–Q4). All maps are plotted on a transparent MNI152 template. The blue to light blue color bars code the thresholded z-statistic value from 3.1 to 6.1.

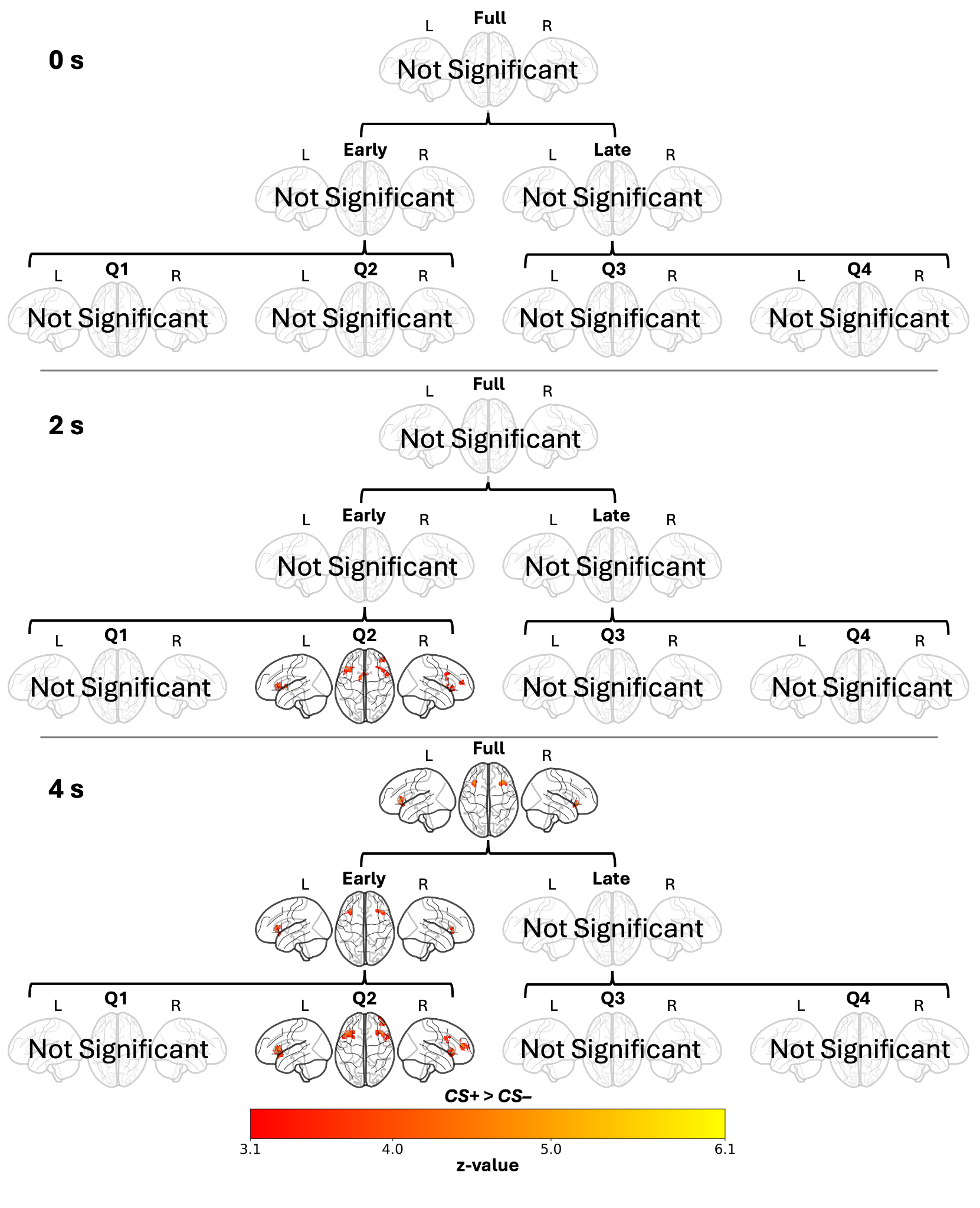

*Figure S3*. Hierarchical BOLD activation patterns for the CS+ > CS– contrast during extinction training, modeled at 0 s, 2 s, and 4 s post-stimulus latencies. The dendrogram structure illustrates the decomposition of the analysis from the Full Training phase (top in each panel), to Halves (middle in each panel), and finally to Quarters (bottom in each panel; Q1–Q4). All maps are plotted on a transparent MNI152 template. The red to yellow color bars code the thresholded z-statistic value from 3.1 to 6.1.

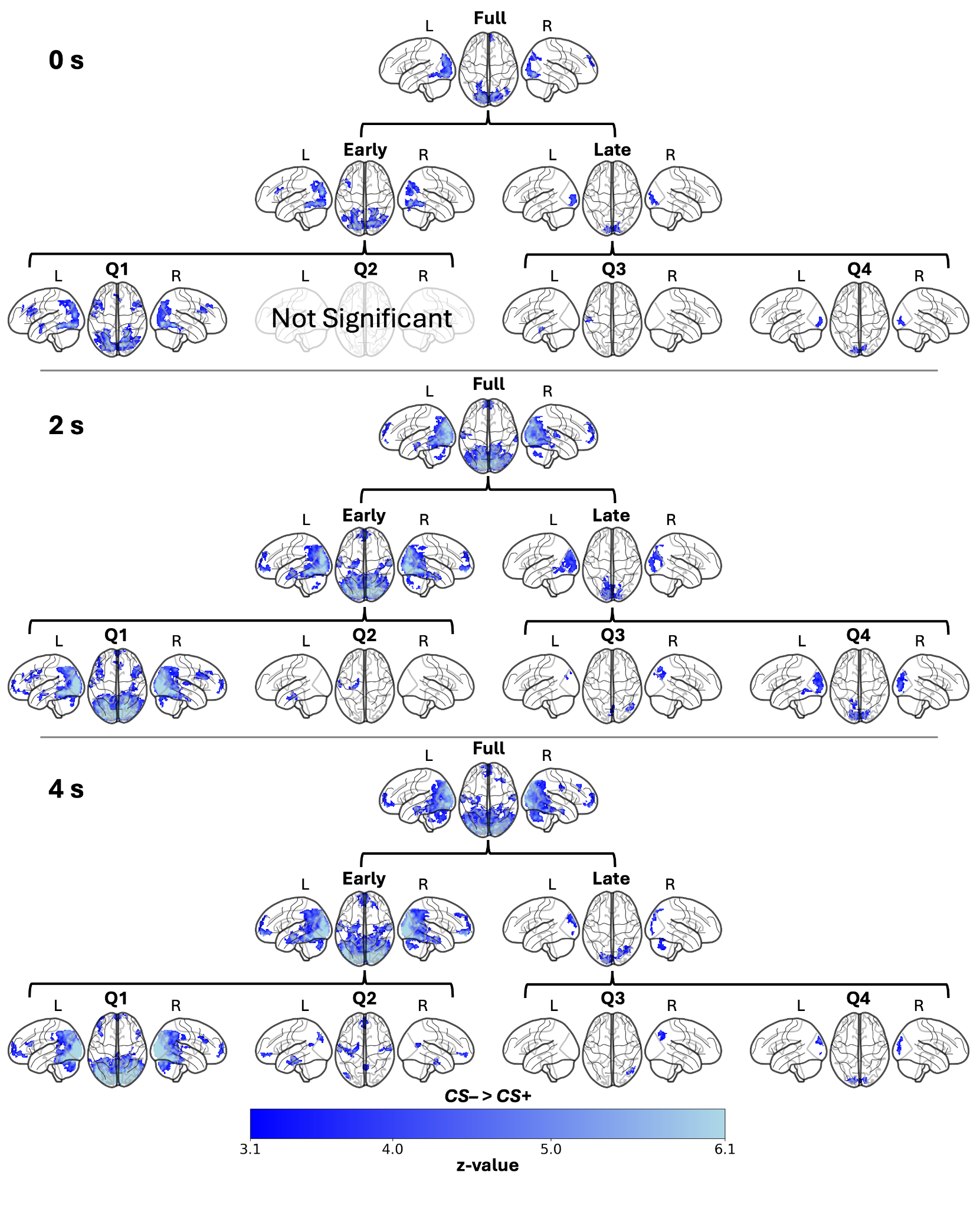

*Figure S4*. Hierarchical BOLD activation patterns for the CS– > CS+ contrast during extinction training, modeled at 0 s, 2 s, and 4 s post-stimulus latencies. The dendrogram structure illustrates the decomposition of the analysis from the Full Training phase (top in each panel), to Halves (middle in each panel), and finally to Quarters (bottom in each panel; Q1–Q4). All maps are plotted on a transparent MNI152 template. The blue to light blue color bars code the thresholded z-statistic value from 3.1 to 6.1.

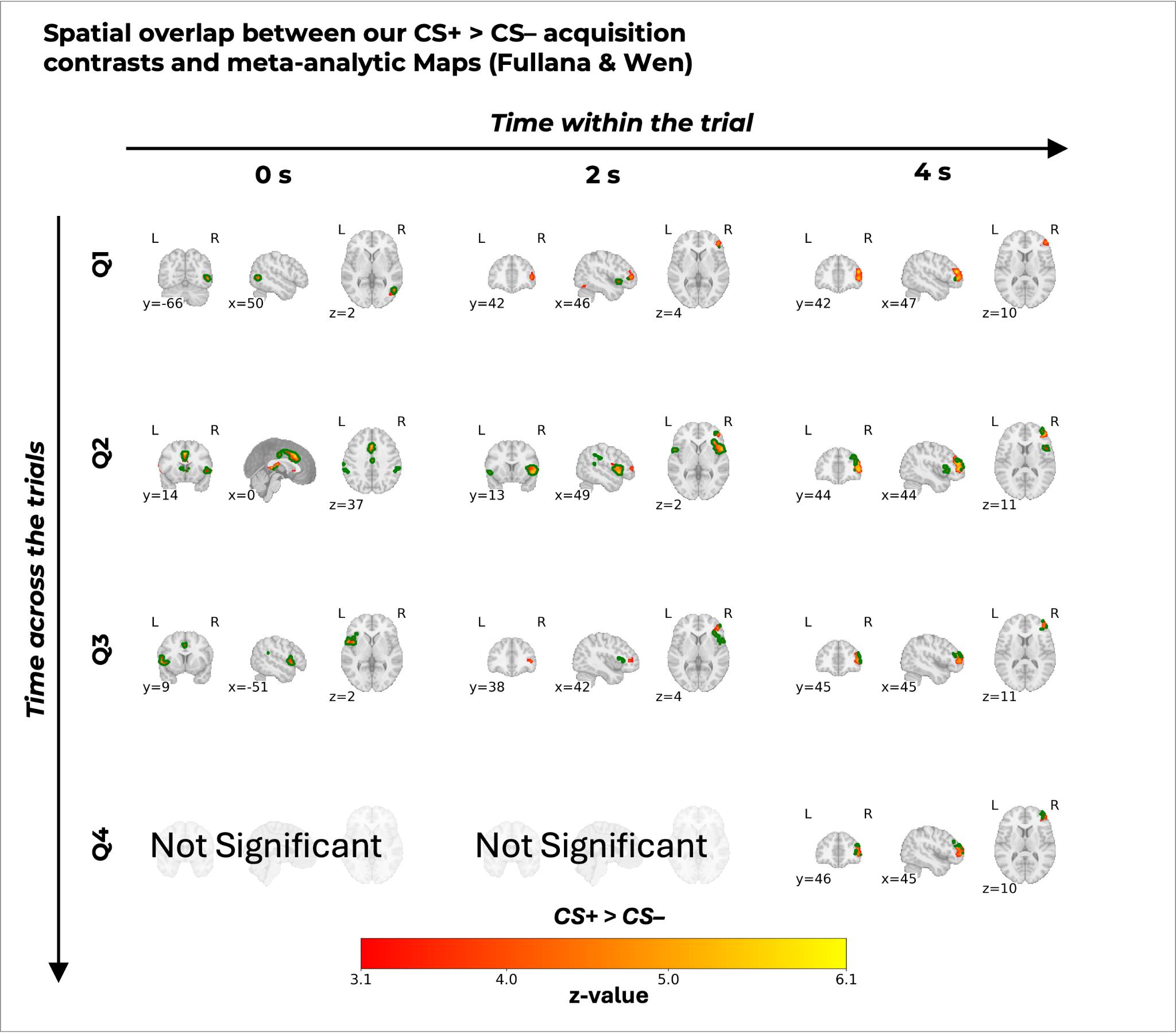

*Figure S5*. Spatial overlap of BOLD activation patterns for the CS+ > CS– contrast (from Figure 3) during fear acquisition training with established fear conditioning networks. The grid illustrates the temporal dynamics of the activations broken down by acquisition quarters (rows; Q1–Q4) and modeled at 0 s, 2 s, and 4 s post-stimulus latencies (columns). Green contours delineate the spatial boundaries of the overlap of significant regions from our analysis and the combined meta-analytic reference maps (Fullana and Wen). Filled, colored regions from red to yellow represent significant clusters from our dataset. "Not Significant" indicates the absence of suprathreshold clusters in our data for that specific time point and quarter. All maps are plotted on a MNI152 template. The colorbar codes the thresholded z-statistic value of our overlaid maps from 3.1 to 6.1.

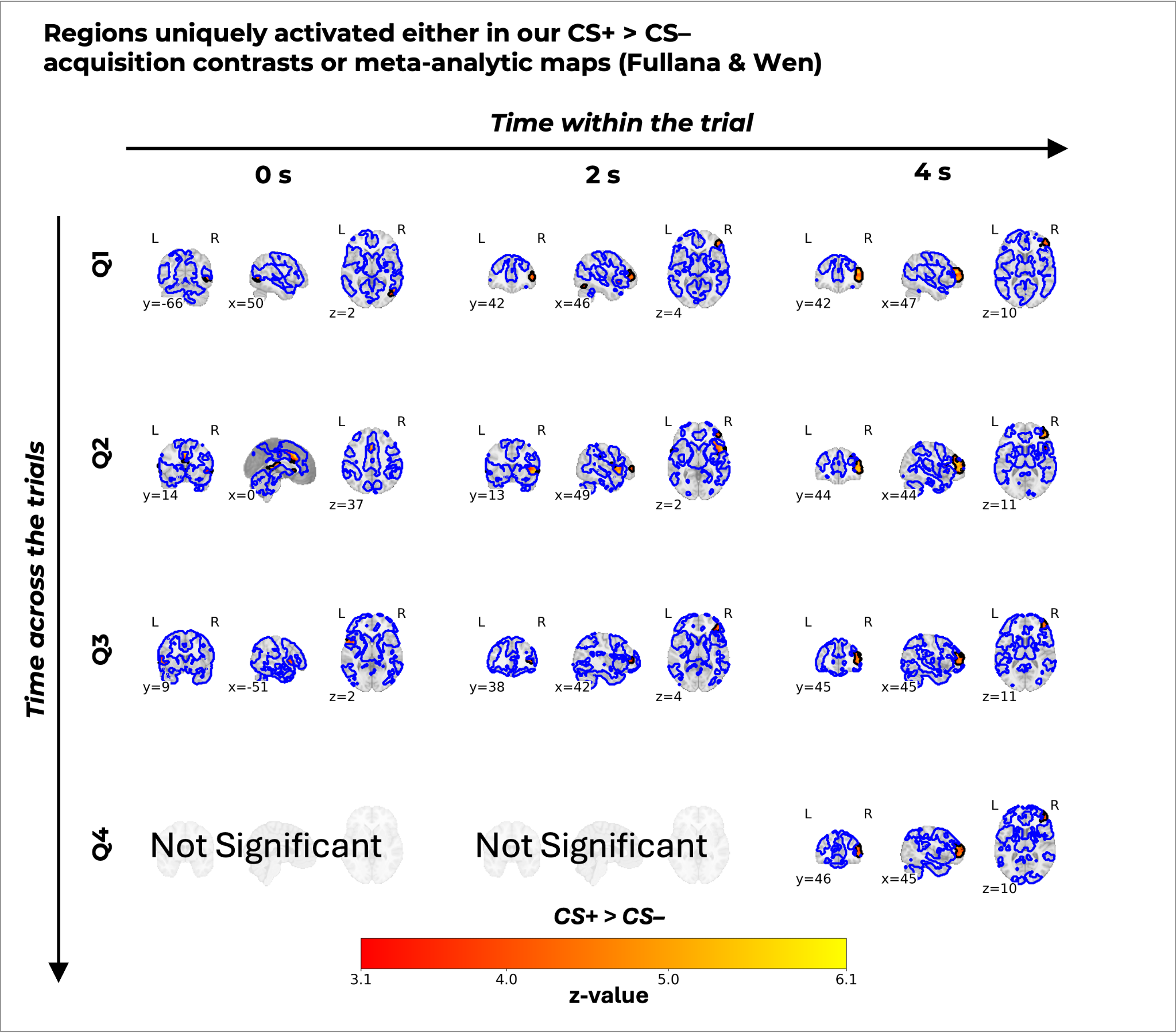

*Figure S6*. Novel BOLD activation patterns for the CS+ > CS– contrast (from Figure 3) during fear acquisition training distinct from established fear conditioning networks. The grid illustrates the temporal dynamics of these unique activations broken down by acquisition quarters (rows; Q1–Q4) and modeled at 0 s, 2 s, and 4 s post-stimulus latencies (columns). Black contours delineate the spatial boundaries of the significant regions uniquely present in our analysis and not in the combined meta-analytic reference maps (Fullana and Wen). Blue contours delineate the spatial boundaries of the significant regions uniquely present in the combined meta-analytic reference maps (Fullana and Wen) and not our analysis. Please note that for CS– > CS+ contrasts in Figures S8, S12, the same contour is presented in red color for improved visibility against the blue to light blue colormap, consistently used in other figures. Filled, colored regions from red to yellow represent significant clusters from our dataset that intersect with these reference networks. "Not Significant" indicates the absence of suprathreshold clusters in our data for that specific time point and quarter. All maps are plotted on an MNI152 template. The colorbar codes the thresholded z-statistic value of our overlaid maps from 3.1 to 6.1.

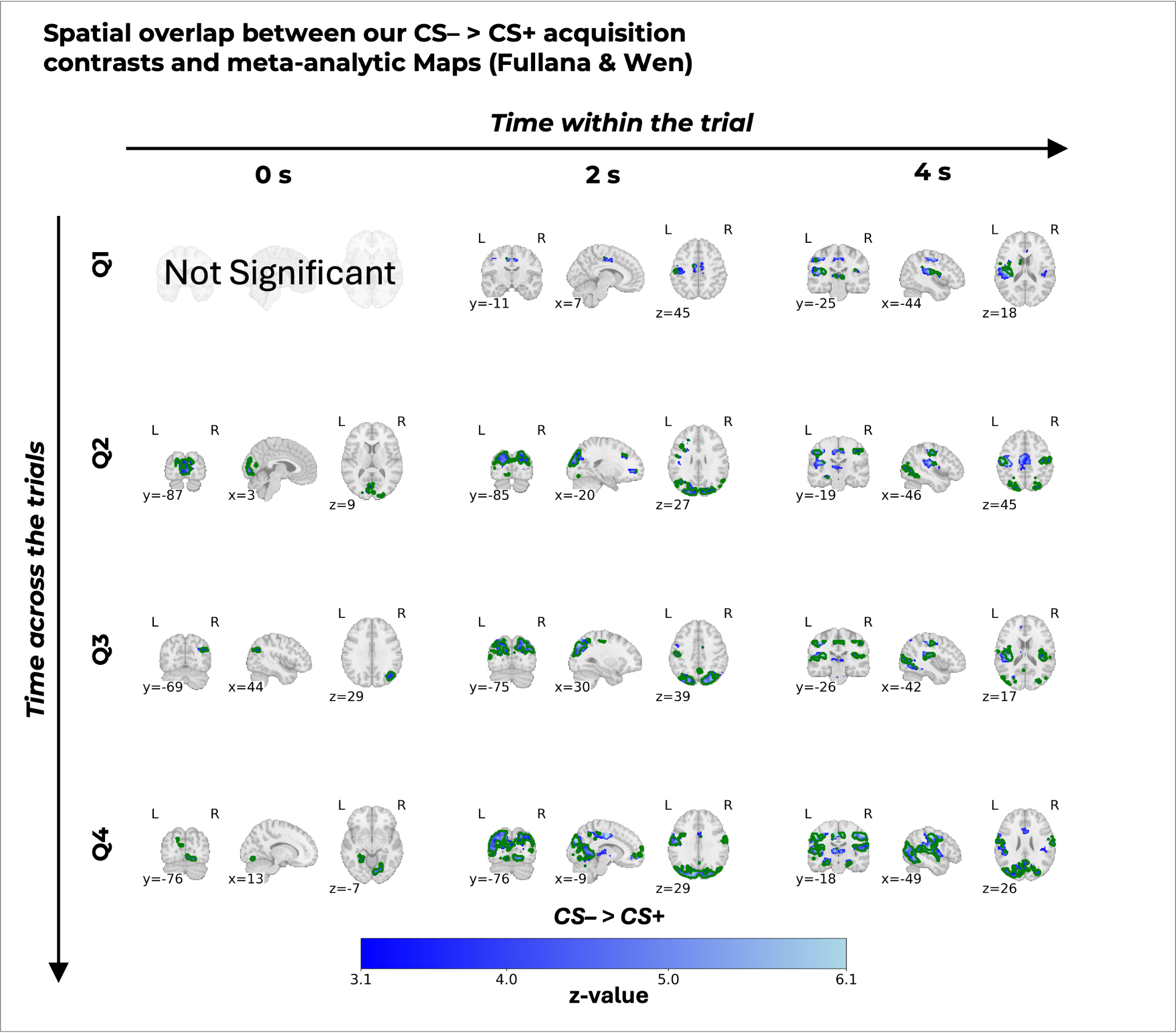

*Figure S7*. Spatial overlap of BOLD activation patterns for the CS– > CS+ contrast (from Figure 3) during fear acquisition training with established fear conditioning networks. The grid illustrates the temporal dynamics of the activations broken down by acquisition quarters (rows; Q1–Q4) and modeled at 0 s, 2 s, and 4 s post-stimulus latencies (columns). Green contours delineate the spatial boundaries of the overlap of significant regions from our analysis and the combined meta-analytic reference maps (Fullana and Wen). Filled, colored regions from blue to light blue represent significant clusters from our dataset. "Not Significant" indicates the absence of suprathreshold clusters in our data for that specific time point and quarter. All maps are plotted on a MNI152 template. The colorbar codes the thresholded z-statistic value of our overlaid maps from 3.1 to 6.1.

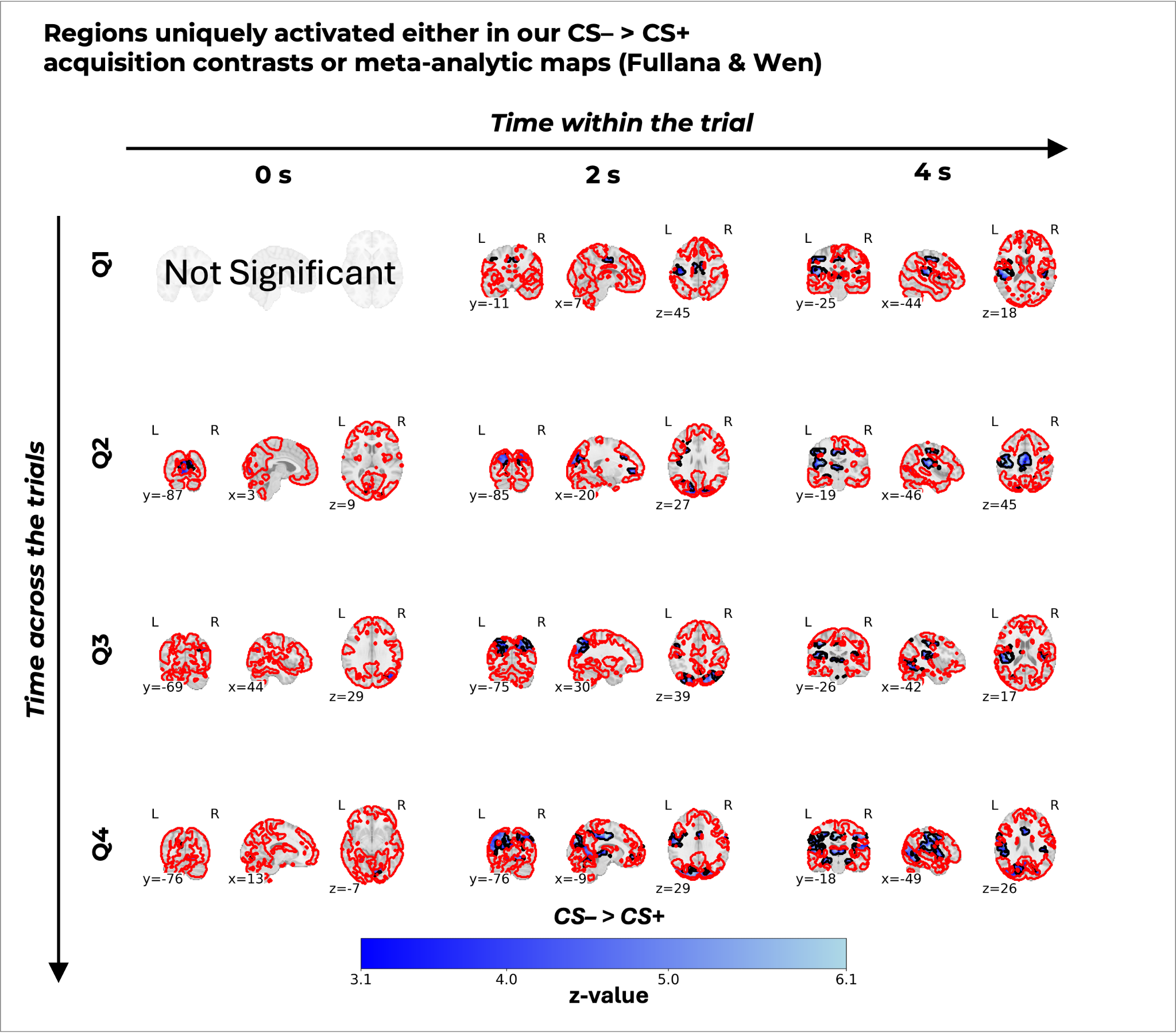

*Figure S8*. Novel BOLD activation patterns for the CS– > CS+ contrast (from Figure 3) during acquisition training distinct from established fear conditioning networks. The grid illustrates the temporal dynamics of these unique activations broken down by acquisition quarters (rows; Q1–Q4) and modeled at 0 s, 2 s, and 4 s post-stimulus latencies (columns). Black contours delineate the spatial boundaries of the significant regions uniquely present in our analysis and not in the combined meta-analytic reference maps (Fullana and Wen). Red contours delineate the spatial boundaries of the significant regions uniquely present in the combined meta-analytic reference maps (Fullana and Wen) and not our analysis. Please note that for CS+ > CS– contrasts in Figures S6, S10, the same contour is presented in blue color for improved visibility against the red to yellow colormap, consistently used in other figures. Filled, colored regions from red to yellow represent significant clusters from our dataset that intersect with these reference networks. "Not Significant" indicates the absence of suprathreshold clusters in our data for that specific time point and quarter. All maps are plotted on an MNI152 template. The colorbar codes the thresholded z-statistic value of our overlaid maps from 3.1 to 6.1.

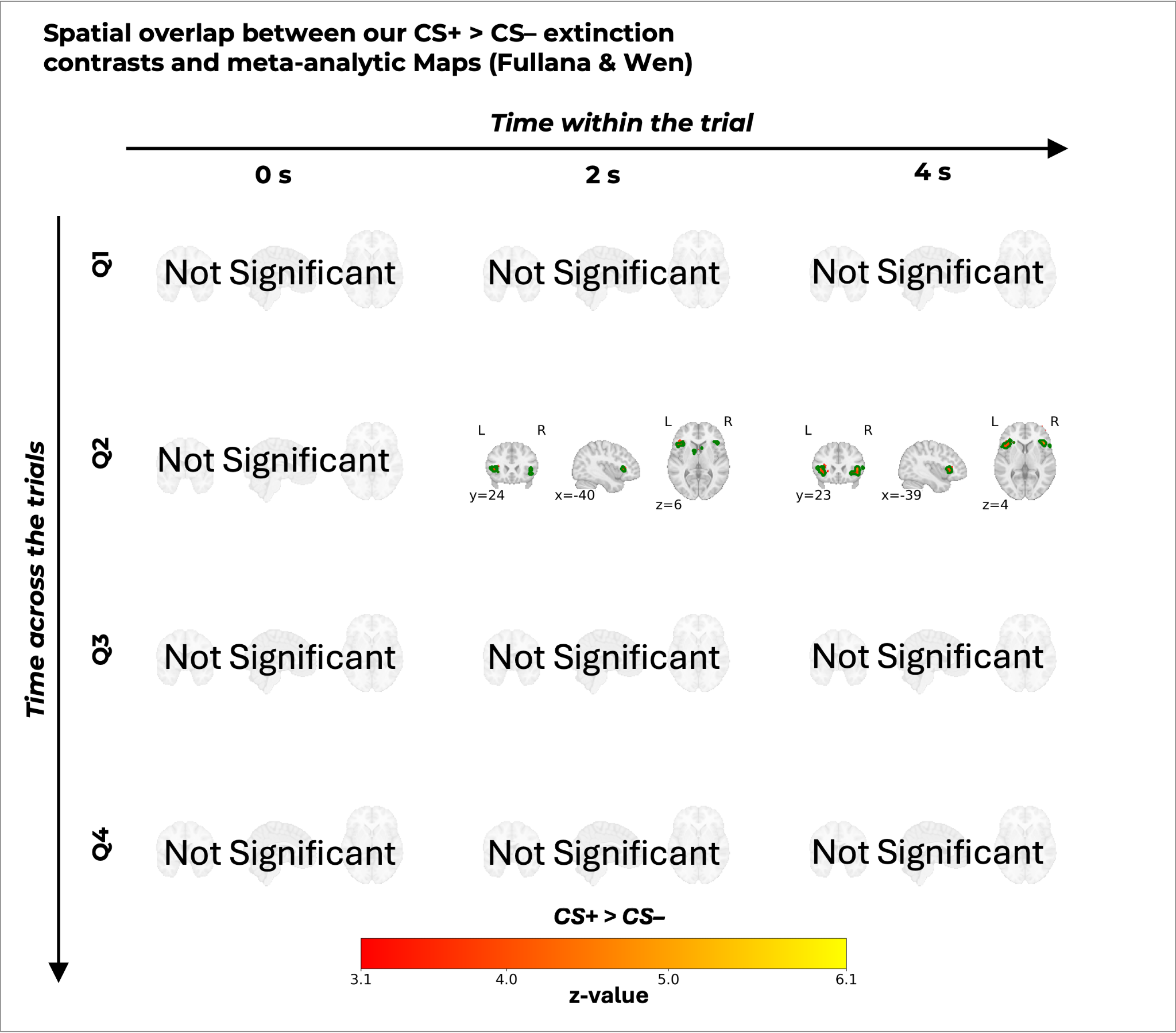

*Figure S9*. Spatial overlap of BOLD activation patterns for the CS+ > CS– contrast (from Figure 4) during extinction training with established extinction networks. The grid illustrates the temporal dynamics of the activations broken down by extinction quarters (rows; Q1–Q4) and modeled at 0 s, 2 s, and 4 s post-stimulus latencies (columns). Green contours delineate the spatial boundaries of the overlap of significant regions from our analysis and the combined meta-analytic reference maps (Fullana and Wen). Filled, colored regions from red to yellow represent significant clusters from our dataset. "Not Significant" indicates the absence of suprathreshold clusters in our data for that specific time point and quarter. All maps are plotted on a MNI152 template. The colorbar codes the thresholded z-statistic value of our overlaid maps from 3.1 to 6.1.

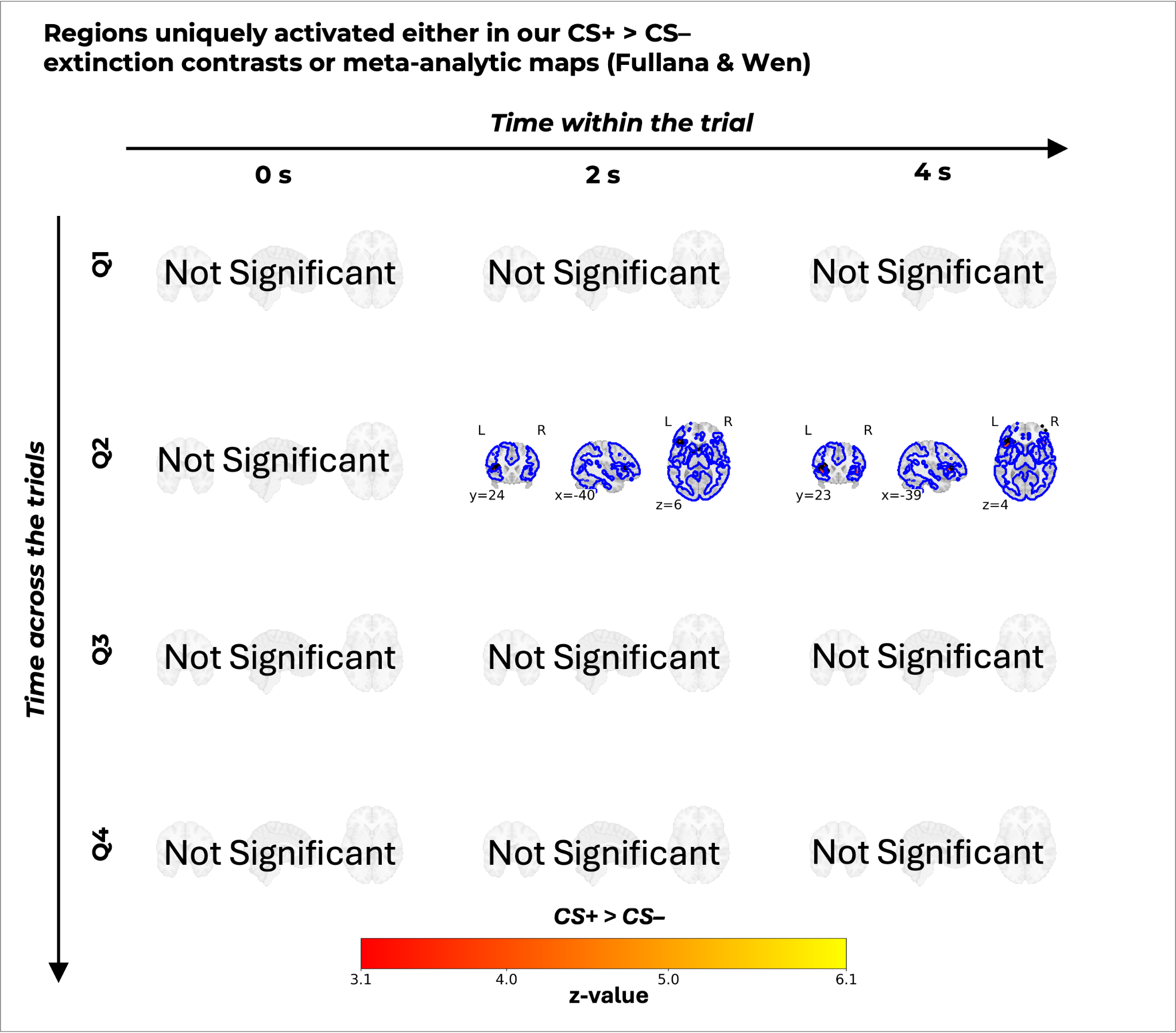

*Figure S10*. Novel BOLD activation patterns for the CS+ > CS– contrast (from Figure 4) during extinction training distinct from established extinction networks. The grid illustrates the temporal dynamics of these unique activations broken down by extinction quarters (rows; Q1–Q4) and modeled at 0 s, 2 s, and 4 s post-stimulus latencies (columns). Black contours delineate the spatial boundaries of the significant regions uniquely present in our analysis and not in the combined meta-analytic reference maps (Fullana and Wen). Blue contours delineate the spatial boundaries of the significant regions uniquely present in the combined meta-analytic reference maps (Fullana and Wen) and not our analysis. Please note that for CS– > CS+ contrasts in Figures S8, S12, the same contour is presented in red color for improved visibility against the blue to light blue colormap, consistently used in other figures. Filled, colored regions from red to yellow represent significant clusters from our dataset that intersect with these reference networks. "Not Significant" indicates the absence of suprathreshold clusters in our data for that specific time point and quarter. All maps are plotted on a MNI152 template. The colorbar codes the thresholded z-statistic value of our overlaid maps from 3.1 to 6.1.

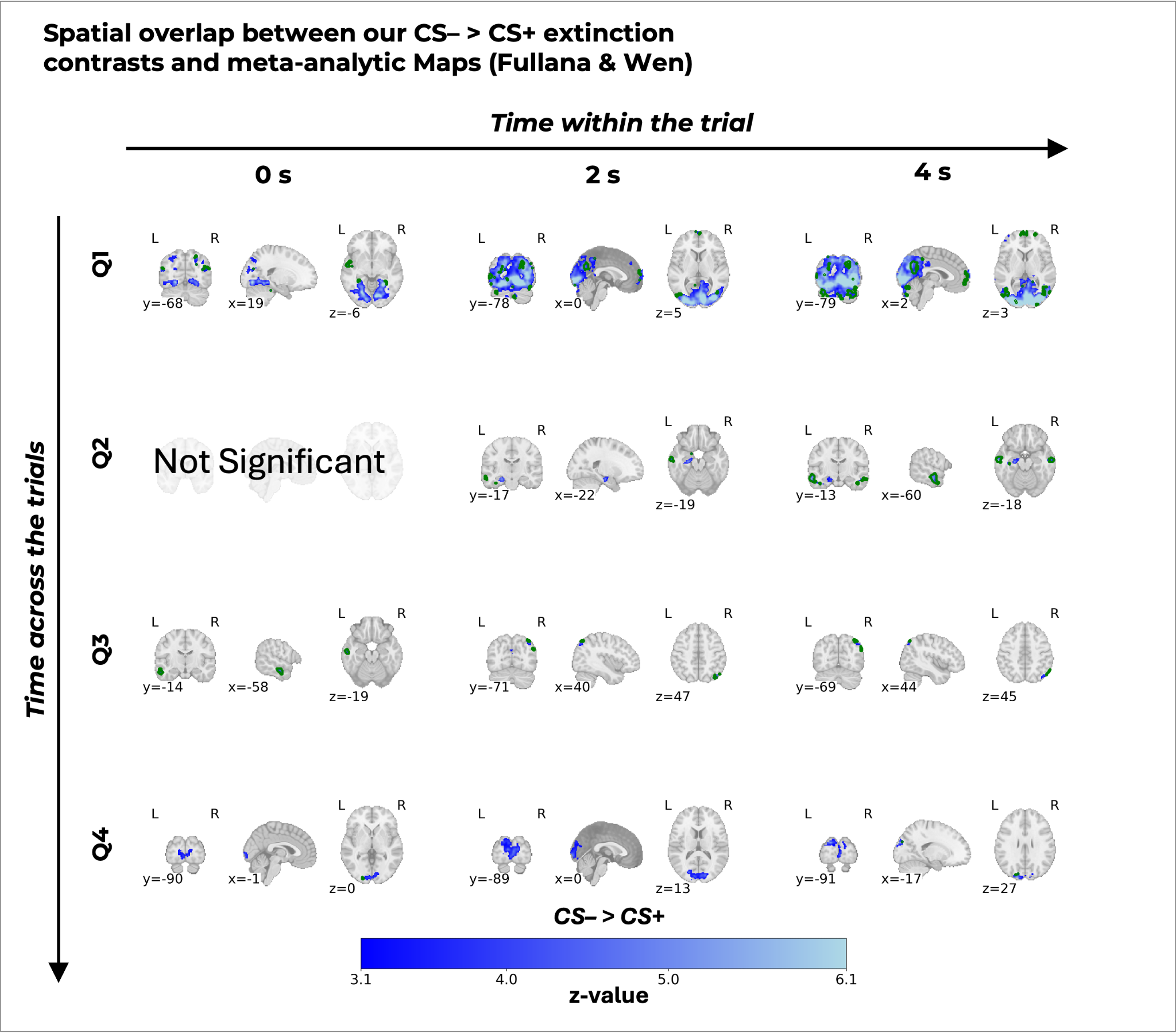

*Figure S11*. Spatial overlap of BOLD activation patterns for the CS– > CS+ contrast (from Figure 4) during extinction training with established extinction networks. The grid illustrates the temporal dynamics of the activations broken down by extinction quarters (rows; Q1–Q4) and modeled at 0 s, 2 s, and 4 s post-stimulus latencies (columns). Green contours delineate the spatial boundaries of the overlap of significant regions from our analysis and the combined meta-analytic reference maps (Fullana and Wen). Filled, colored regions from blue to light blue represent significant clusters from our dataset. "Not Significant" indicates the absence of suprathreshold clusters in our data for that specific time point and quarter. All maps are plotted on a MNI152 template. The colorbar codes the thresholded z-statistic value of our overlaid maps from 3.1 to 6.1.

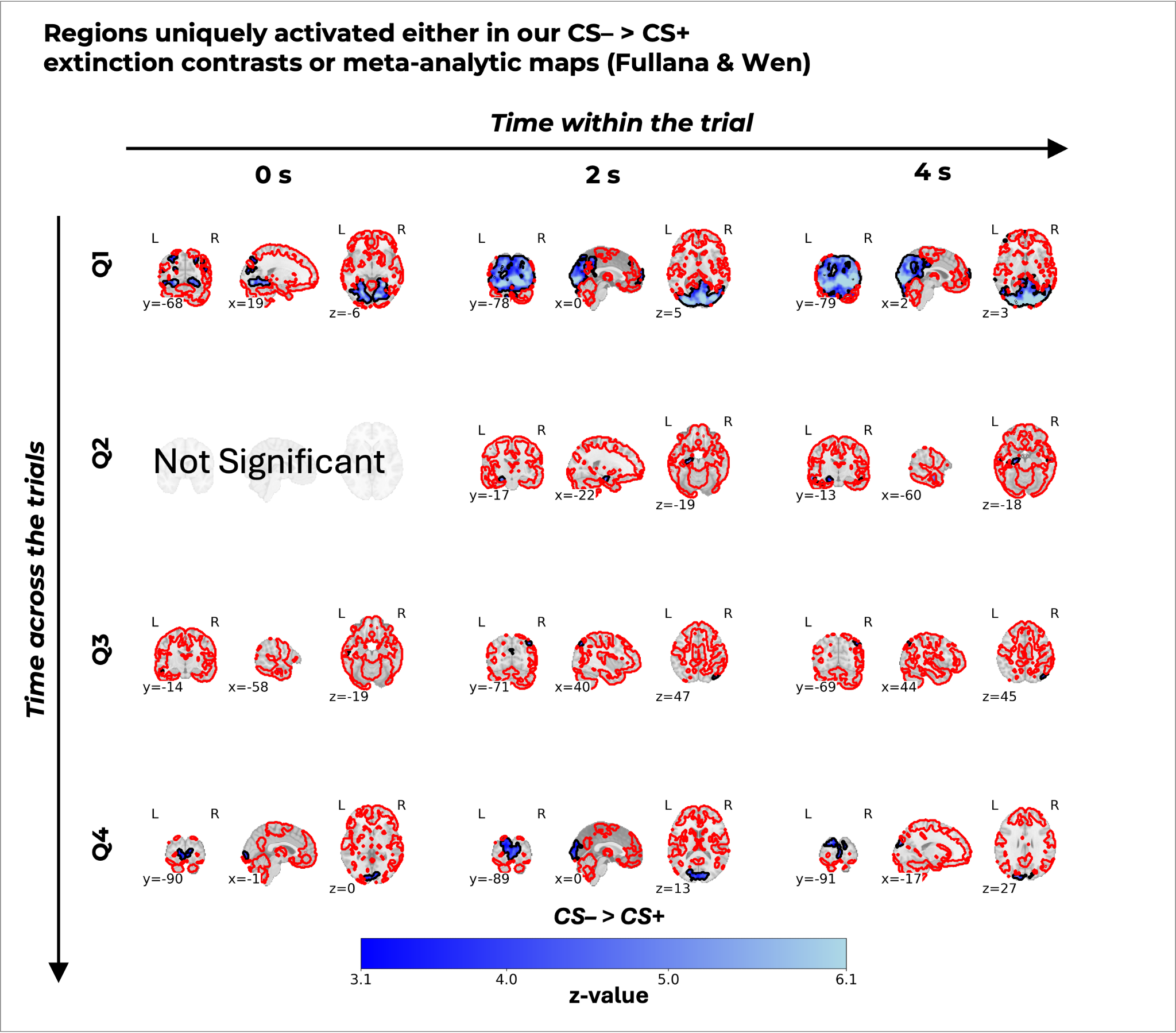

*Figure S12*. Novel BOLD activation patterns for the CS– > CS+ contrast (from Figure 4) during extinction training distinct from established extinction networks. The grid illustrates the temporal dynamics of these unique activations broken down by extinction quarters (rows; Q1–Q4) and modeled at 0 s, 2 s, and 4 s post-stimulus latencies (columns). Black contours delineate the spatial boundaries of the significant regions uniquely present in our analysis and not in the combined meta-analytic reference maps (Fullana and Wen). Red contours delineate the spatial boundaries of the significant regions uniquely present in the combined meta-analytic reference maps (Fullana and Wen) and not our analysis. Please note that for CS+ > CS– contrasts in Figures S6, S10, the same contour is presented in blue color for improved visibility against the red to yellow colormap, consistently used in other figures. Filled, colored regions from red to yellow represent significant clusters from our dataset that intersect with these reference networks. "Not Significant" indicates the absence of suprathreshold clusters in our data for that specific time point and quarter. All maps are plotted on a MNI152 template. The colorbar codes the thresholded z-statistic value of our overlaid maps from 3.1 to 6.1.

Table S1. Overview of significant anatomical areas from the full analysis (average of all trials in respective training phase) during fear acquisition and extinction training. dACC = dorsal anterior cingulate cortex; PCC = posterior cingulate cortex; vmPFC = ventromedial prefrontal cortex; vlPFC = ventrolateral prefrontal cortex; dlPFC = dorsolateral prefrontal cortex.

| **Phase** | **Time** | **Contrast** | **Area** |
| --- | --- | --- | --- |
| Acquisition | 0s | CS+ > CS− | *dACC, PCC, anterior and posterior callosal body, fornix, visual cortex V4-V5, insular cortex, left insular cortex, bilateral operculum* |
| Acquisition | 0s | CS− > CS+ | *Left parahippocampal, bilateral visual cortex V1-V4, right inferior parietal lobule, left superior parietal lobule* |
| Acquisition | 2s | CS+ > CS− | *Bilateral visual cortex V4, left parietal operculum, left Broca's area, left frontal operculum, left precentral gyrus, left secondary somatosensory, left frontal pole, right frontal operculum* |
| Acquisition | 2s | CS− > CS+ | *Major activations across the brain, including bilateral vmpfc, bilateral cuneus, left hippocampus and parahippocampus, bilateral operculum, bilateral inferior parietal lobule* |
| Acquisition | 4s | CS+ > CS− | *Left frontal operculum, left frontal pole* |
| Acquisition | 4s | CS− > CS+ | *Major activations across the brain, including dACC, bilateral primary motor cortex, bilateral premotor cortex, and bilateral vmPFC* |
| Extinction | 0s | CS+ > CS− | *No significant activation* |
| Extinction | 0s | CS− > CS+ | *Bilateral visual cortex V1-V4, right dmPFC as part of the superior frontal gyrus* |
| Extinction | 2s | CS+ > CS− | *No significant activation* |
| Extinction | 2s | CS− > CS+ | *Bilateral visual cortex V1-V4, bilateral cuneus and precuneus cortices, bilateral dmPFC, bilateral hippocampus and parahippocampus* |
| Extinction | 4s | CS+ > CS− | *Bilateral frontal operculum and insula* |
| Extinction | 4s | CS− > CS+ | *Bilateral visual cortex V1-V4, bilateral cuneus and precuneus cortices, Left amygdala, bilateral hippocampus (stronger left activation), right primary somatosensory and motor cortices, bilateral dmPFC and vmPFC* |

Table S2. Overview of significant anatomical areas split into halves and trial segments during fear acquisition training. dACC = dorsal anterior cingulate cortex; PCC = posterior cingulate cortex; vmPFC = ventromedial prefrontal cortex; vlPFC = ventrolateral prefrontal cortex; dlPFC = dorsolateral prefrontal cortex.

| **Phase** | **Time** | **Contrast** | **Half** | **Area** |
| --- | --- | --- | --- | --- |
| Acquisition | 0s | CS+ > CS− | *Early* | *dACC, PCC, fornix extending to bilateral thalamus, bilateral secondary somatosensory cortex and central operculum, bilateral parietal operculum* |
| Acquisition | 0s | CS+ > CS− | *Late* | *dACC, left operculum* |
| Acquisition | 0s | CS− > CS+ | *Early* | *Right inferior parietal lobule* |
| Acquisition | 0s | CS− > CS+ | *Late* | *Left hippocampus, left parahippocampus, visual cortex (V2-V4), left cuneus and precuneus, right inferior parietal lobule, right insula and operculum* |
| Acquisition | 2s | CS+ > CS− | *Early* | *Right central operculum, frontal operculum, and insula (contiguous cluster), bilateral visual cortex (V1-V4), right vlPFC, right primary somatosensory cortex extending to inferior-anterior parietal lobule* |
| Acquisition | 2s | CS+ > CS− | *Late* | *Right VLPFC, right insula and frontal operculum* |
| Acquisition | 2s | CS− > CS+ | *Early* | *Right inferior parietal lobule, bilateral visual cortex (V1-V4), bilateral premotor cortex, cingulate gyrus, and primary motor cortex, left insula and operculum, left primary somatosensory cortex, left vmPFC, left vlPFC* |
| Acquisition | 2s | CS− > CS+ | *Late* | *Bilateral hippocampus, parahippocampus, amygdala, visual cortex (V1-V4), bilateral cuneus and precuneus, thalamus, brainstem, vmPFC, bilateral premotor cortex, bilateral operculum, bilateral insula, cerebellar vermis and crus (multiple lobules)* |
| Acquisition | 4s | CS+ > CS− | *Early* | *Right frontal operculum and insula, right vlPFC* |
| Acquisition | 4s | CS+ > CS− | *Late* | *Right vlPFC* |
| Acquisition | 4s | CS− > CS+ | *Early* | *Bilateral premotor cortex, primary and secondary somatosensory cortex, left inferior parietal lobule, left V5, bilateral vmPFC, left hippocampus, thalamus, and fornix* |
| Acquisition | 4s | CS− > CS+ | *Late* | *Bilateral premotor cortex, primary and secondary somatosensory cortex, left inferior parietal lobule, left V5, bilateral vmPFC, left hippocampus, thalamus, fornix, amygdala* |
| Extinction | 0s | CS+ > CS− | *Early* | *No significant activation* |
| Extinction | 0s | CS+ > CS− | *Late* | *No significant activation* |
| Extinction | 0s | CS− > CS+ | *Early* | *Bilateral visual cortex (V1-V4), right cerebellar crus I and II, vermis VI, left middle frontal gyrus* |
| Extinction | 0s | CS− > CS+ | *Late* | *Bilateral visual cortex (V1-V4)* |
| Extinction | 2s | CS+ > CS− | *Early* | *No significant activation* |
| Extinction | 2s | CS+ > CS− | *Late* | *No significant activation* |
| Extinction | 2s | CS− > CS+ | *Early* | *Extensive bilateral visual cortex, cuneus, and precuneus, left amygdala, bilateral hippocampus, vmPFC* |
| Extinction | 2s | CS− > CS+ | *Late* | *Bilateral visual cortex (V1-V4), bilateral cuneus, left precuneus, left hippocampus* |
| Extinction | 4s | CS+ > CS− | *Early* | *Bilateral frontal operculum and insula* |
| Extinction | 4s | CS+ > CS− | *Late* | *No significant activation* |
| Extinction | 4s | CS− > CS+ | *Early* | *Bilateral hippocampus and parahippocampus, left amygdala, bilateral cuneus and precuneus, vmPFC, dmPFC, bilateral visual cortex (V1-V5), cerebellum (right lobules I-IV and V, bilateral crus I and V, right vermis crus II)* |
| Extinction | 4s | CS− > CS+ | *Late* | *Bilateral cuneus, right precuneus and superior parietal lobule, cerebellum (right crus I and II, right lobules VI and VIIb)* |

Table S3. Overview of significant anatomical areas split into quarters and trial segments during fear acquisition training. dACC = dorsal anterior cingulate cortex; PCC = posterior cingulate cortex; vmPFC = ventromedial prefrontal cortex; vlPFC = ventrolateral prefrontal cortex; dlPFC = dorsolateral prefrontal cortex.

| **Time** | **Contrast** | **Quarter** | **Area** |
| --- | --- | --- | --- |
| 0s | CS+ > CS− | *Q1* | *Right Visual Cortex (V4 and V5)* |
| 0s | CS+ > CS− | *Q2* | *dACC, cingulum, callosal body, bilateral parietal operculum extending to secondary somatosensory cortex (S2), fornix extending to bilateral hippocampus (cornu ammonis and dentate gyrus), frontal operculum, left primary somatosensory cortex (S1), right putamen, right cerebellar crus I, and left caudate* |
| 0s | CS+ > CS− | *Q3* | *dACC, left secondary somatosensory cortex and parietal operculum (OP4) extending to primary somatosensory cortex and precentral gyrus, frontal operculum* |
| 0s | CS+ > CS− | *Q4* | *No significant activation* |
| 0s | CS− > CS+ | *Q1* | *No significant activation* |
| 0s | CS− > CS+ | *Q2* | *Left visual cortex (V1-V4) and left cuneal cortex* |
| 0s | CS− > CS+ | *Q3* | *Right inferior parietal lobule, left visual cortex (V2-V4)* |
| 0s | CS− > CS+ | *Q4* | *Left precuneal and cuneal cortices, left hippocampus extending to parahippocampal gyrus and cingulum, bilateral secondary somatosensory cortex extending to right insula* |
| 2s | CS+ > CS− | *Q1* | *Right frontal operculum, right vlPFC, right visual cortex (V4-V5)* |
| 2s | CS+ > CS− | *Q2* | *Right dlPFC, right insula, right parietal operculum extending to right frontal operculum (contiguous cluster), right somatosensory cortex, bilateral inferior parietal lobule* |
| 2s | CS+ > CS− | *Q3* | *Right supramarginal gyrus extending to right inferior parietal lobule, left inferior parietal lobule, right somatosensory cortex, right frontal operculum, right dlPFC* |
| 2s | CS+ > CS− | *Q4* | *No significant activation* |
| 2s | CS− > CS+ | *Q1* | *No significant activation* |
| 2s | CS− > CS+ | *Q2* | *Bilateral visual cortex (V1-V4)* |
| 2s | CS− > CS+ | *Q3* | *Left visual cortex (V1-V4), right inferior parietal lobule* |
| 2s | CS− > CS+ | *Q4* | *Left hippocampus extending to parahippocampal gyrus and cingulum, bilateral parietal operculum, right secondary somatosensory cortex, right visual cortex (V1-V4), left cuneal cortex, left cerebellar lobules I-IV and V* |
| 4s | CS+ > CS− | *Q1* | *Right vlPFC* |
| 4s | CS+ > CS− | *Q2* | *Right vlPFC, right frontal operculum* |
| 4s | CS+ > CS− | *Q3* | *Right vlPFC* |
| 4s | CS+ > CS− | *Q4* | *Right vlPFC* |
| 4s | CS− > CS+ | *Q1* | *Widespread activation across multiple regions, pattern overlaps primarily with Fullana et al. fear and safety networks* |
| 4s | CS− > CS+ | *Q2* | *Widespread activation across multiple regions, pattern overlaps primarily with Fullana et al. fear and safety networks* |
| 4s | CS− > CS+ | *Q3* | *Widespread activation across multiple regions, pattern overlaps primarily with Fullana et al. fear and safety networks* |
| 4s | CS− > CS+ | *Q4* | *Widespread activation across multiple regions, pattern overlaps primarily with Fullana et al. fear and safety networks* |

Table S4. Overview of significant anatomical areas split into quarters and trial segments during fear extinction training. dACC = dorsal anterior cingulate cortex; PCC = posterior cingulate cortex; vmPFC = ventromedial prefrontal cortex; vlPFC = ventrolateral prefrontal cortex; dlPFC = dorsolateral prefrontal cortex.

| **Time** | **Contrast** | **Quarter** | **Area** |
| --- | --- | --- | --- |
| 0s | CS+ > CS− | *Q1* | *No significant activation* |
| 0s | CS+ > CS− | *Q2* | *No significant activation* |
| 0s | CS+ > CS− | *Q3* | *No significant activation* |
| 0s | CS+ > CS− | *Q4* | *No significant activation* |
| 0s | CS− > CS+ | *Q1* | *Bilateral visual cortex (V1-V4), right posterior parahippocampus, bilateral cuneal and precuneal cortices extending to bilateral inferior and superior parietal lobules, bilateral frontal operculum, paracingulate cortex* |
| 0s | CS− > CS+ | *Q2* | *No significant activation* |
| 0s | CS− > CS+ | *Q3* | *Left middle temporal gyrus* |
| 0s | CS− > CS+ | *Q4* | *Bilateral visual cortex (V1-V4)* |
| 2s | CS+ > CS− | *Q1* | *No significant activation* |
| 2s | CS+ > CS− | *Q2* | *Bilateral insula (left predominant), bilateral operculum (left predominant), left thalamus extending to right caudate, dorsolateral prefrontal cortex (dlPFC)* |
| 2s | CS+ > CS− | *Q3* | *No significant activation* |
| 2s | CS+ > CS− | *Q4* | *No significant activation* |
| 2s | CS− > CS+ | *Q1* | *Visual cortex, bilateral hippocampus, bilateral parahippocampus, paracingulate cortex, right precentral gyrus extending to right inferior frontal gyrus, right insula, bilateral inferior parietal lobule, vmPFC* |
| 2s | CS− > CS+ | *Q2* | *Left hippocampus, left middle temporal gyrus* |
| 2s | CS− > CS+ | *Q3* | *Right superior parietal lobule, central cuneal and precuneal cortices, right inferior parietal lobule* |
| 2s | CS− > CS+ | *Q4* | *Visual cortex (V1-V4, left predominant), left cuneus* |
| 4s | CS+ > CS− | *Q1* | *No significant activation* |
| 4s | CS+ > CS− | *Q2* | *Left frontal operculum extending to left insula, dlPFC, right frontal operculum* |
| 4s | CS+ > CS− | *Q3* | *No significant activation* |
| 4s | CS+ > CS− | *Q4* | *No significant activation* |
| 4s | CS− > CS+ | *Q1* | *Extensive bilateral visual cortex (V1-V4), bilateral cerebellar crus I and II, right cerebellar lobules V and I-IV, posterior cingulate cortex, vmPFC, left dlPFC, right inferior frontal gyrus, left frontal pole* |
| 4s | CS− > CS+ | *Q2* | *Left amygdala, left parahippocampus, bilateral hippocampus, bilateral middle temporal gyrus, ventromedial prefrontal cortex (vmPFC), left inferior parietal lobule, cingulum, bilateral cuneal and precuneal cortices* |
| 4s | CS− > CS+ | *Q3* | *Right inferior parietal lobule, angular gyrus* |
| 4s | CS− > CS+ | *Q4* | *Visual cortex (V1-V2), left cuneus* |
